# TRAIL promotes the polarization of human macrophages toward a proinflammatory M1 phenotype and is associated with increased survival in cancer patients with high tumor macrophage content

**DOI:** 10.1101/2023.08.16.553509

**Authors:** Sinem Gunalp, Derya Goksu Helvacı, Aysenur Oner, Ahmet Bursalı, Alessandra Conforte, Hüseyin Güner, Gökhan Karakülah, Eva Szegezdi, Duygu Sag

## Abstract

**Background:** TNF-related apoptosis-inducing ligand (TRAIL) is a member of the TNF superfamily that can either induce cell death or activate survival pathways after binding to death receptors (DRs) DR4 or DR5. TRAIL is investigated as a therapeutic agent in clinical trials due to its selective toxicity to transformed cells.

Macrophages can be polarized into pro-inflammatory/tumor-fighting M1 macrophages or anti-inflammatory/tumor-supportive M2 macrophages and an inbalance between M1 and M2 macrophages can promote diseases. Therefore, identifying modulators that regulate macrophage polarization is important to design effective macrophage-targeted immunotherapies. The impact of TRAIL on macrophage polarization is not known.

**Methods:** Primary human monocyte-derived macrophages were pre-treated with either TRAIL or with DR4 or DR5-specific ligands and then polarized into M1, M2a, or M2c phenotypes *in vitro*. The expression of M1 and M2 markers in macrophage subtypes was analyzed by RNA sequencing, qPCR, ELISA, and flow cytometry. Furthermore, the cytotoxicity of the macrophages against U937 AML tumor targets was assessed by flow cytometry. TCGA datasets were also analyzed to correlate TRAIL with M1/M2 markers, and the overall survival of cancer patients.

**Results:** TRAIL increased the expression of M1 markers at both mRNA and protein levels while decreasing the expression of M2 markers at the mRNA level in human macrophages. TRAIL also shifted M2 macrophages towards an M1 phenotype. Our data showed that both DR4 and DR5 death receptors play a role in macrophage polarization. Furthermore, TRAIL enhanced the cytotoxicity of macrophages against the AML cancer cells *in vitro*. Finally, TRAIL expression was positively correlated with increased expression of M1 markers in the tumors from ovarian and sarcoma cancer patients and longer overall survival in cases with high, but not low, tumor macrophage content.

**Conclusions:** TRAIL promotes the polarization of human macrophages toward a proinflammatory M1 phenotype via both DR4 and DR5. Our study defines TRAIL as a new regulator of macrophage polarization and suggests that targeting DRs can enhance the anti-tumorigenic response of macrophages in the tumor microenvironment by increasing M1 polarization.

## INTRODUCTION

Macrophages are myeloid-origin immune cells that play key roles in host defense, homeostasis and immune surveillance (1, 2). They can be polarized into pro-inflammatory M1 or anti-inflammatory M2 macrophages in the presence of certain polarization factors (3, 4). LPS/IFN-γ, or TNFα stimulation drives macrophages into an M1 phenotype (5, 6). M2 macrophages are further subdivided into M2a, M2b and M2c phenotypes. While IL-4 and/or IL-13 induce M2a polarization; immune complex and TLR/IL-1R ligands drive macrophages into M2b; and IL-10, TGFβ or glucocorticoids polarize macrophages into M2c phenotypes (5, 6, 7, 8). While M1 macrophages play a key role in host defense against pathogen invasion and tumor-fighting, M2 macrophages, according to their subgroups, are involved in allergic reactions, tissue regeneration, and tumor growth (5, 9, 10). Macrophage polarization is highly dynamic (1) and the loss of M1/M2 balance can form the basis of various diseases including obesity, type 2 diabetes, viral infections, allergic asthma, and cancer (11, 12, 13, 14, 15, 16, 17, 18, 19). Therefore, elucidating the mechanisms that regulate macrophage polarization is crucial for finding possible treatments for these diseases.

TNF-related apoptosis-inducing ligand (TRAIL), as TNF and Fas ligand, is a member of the TNF superfamily (20, 21, 22, 23). In human cells, TRAIL can bind to the death receptors DR4, and DR5, which induces cell death or survival pathways (22, 24, 25, 26); or to the decoy receptors; DcR1, DcR2, and OPG, which can neutralize the activation of DR4 and DR5 (27, 28, 29, 30, 31). The state of the cell, such as the ratio of anti-apoptotic/apoptotic molecules, or the distribution of functional and decoy receptors could determine which pathway to proceed upon DR4 and DR5 activation (32, 33, 34, 35, 36, 37). In the target cell, TRAIL can trigger extrinsic/intrinsic apoptotic pathways (38, 39, 40) or necroptosis (24, 41). Besides cell death, survival pathways involving migration, proliferation, and inflammatory cytokine production can also be induced (22, 26).

Transformed cells are more susceptible to TRAIL-induced cell death than normal cells which led to the widespread study of TRAIL in cancer therapies (22, 24). TRAIL can prevent tumor growth and metastasis in tumor-bearing mouse models (42, 43, 44) and induce cell death in tumor-promoting cells including tumor-associated M2 type macrophages (TAMs) and endothelial cells (45, 46, 47, 48). For TRAIL-resistant tumors (49, 50, 51) various formulations (52, 53) and tumor-specific approaches are being developed to increase the efficacy of DR4/5 activation (54, 55, 56, 57).

Human monocyte-derived macrophages express both functional; DR4 and DR5, and decoy; DcR1 and DcR2 death receptors (34, 58, 59). Studies to date mostly have covered the sensitivity of macrophage subtypes to TRAIL in a context-dependent manner. For instance, while M1 macrophages are more susceptible to TRAIL-induced cell death in autoimmune diseases (60, 61), this scenario applies to M2 macrophages in the tumor microenvironment (TME) (34, 47, 48, 62). Additionally, studies demonstrated that TRAIL can increase the expression of inflammatory cytokines from primary human/mouse macrophages (63, 64) and also from tumor-associated macrophages (TAMs) in tumor-bearing mice (63). Contrary to this, another study reported that TRAIL can restrain inflammation-induced tumor formation by decreasing the number of inflammatory macrophages in the target tissue (65). However, the role of TRAIL in human macrophage polarization and function is not known.

In this study, we demonstrate that TRAIL polarizes primary human macrophages into a pro-inflammatory M1 phenotype by increasing the expression of M1 markers while decreasing the expression of M2 markers in macrophage subtypes through both DR4 and DR5 death receptors. TRAIL enhances macrophage cytotoxicity against cancer cells and high TRAIL expression in the TME is positively correlated with the expression of M1 markers in cancer patients and longer overall survival in cases with high, but not low, tumor macrophage content.

## MATERIALS and METHODS

### Study approval

Buffy coats of healthy donors were obtained from Dokuz Eylul University Blood Bank (Izmir, Turkey) and University of Galway (Galway, Ireland). Ethical approval was provided by the Non-Interventional Research Ethics Committee of Dokuz Eylul University (Approval number: 2018/06-26) and University of Galway Research Ethics Committee (Approval number: 2022.02.022).

### Generation of primary human monocyte-derived macrophages

Primary human monocytes were isolated from buffy coats of healthy donors between the ages of 25 and 45. First, by density gradient centrifugation technique, peripheral blood mononuclear cells (PBMCs) were isolated from the buffy coats with Ficoll-Paque (GE Healthcare, Pittsburgh, PA). Subsequently, monocytes were isolated from PBMCs with Percoll (GE Healthcare, Pittsburgh, PA) by a second density gradient centrifugation (66).

Isolated monocytes were cultured in R5 medium [RPMI 1640 (Gibco, ThermoFisher Scientific, Waltham, MA, USA) supplemented with 5% heat-inactivated fetal bovine serum (Gibco, ThermoFisher Scientific, Waltham, MA, USA), 1% L-glutamine (Gibco, ThermoFisher Scientific, Waltham, MA, USA) and 1% Penicillin-Streptomycin (Gibco, ThermoFisher Scientific, Waltham, MA, USA)] in the presence of 10 ng/mL human recombinant M-CSF (PeproTech, Rocky Hill, NJ). Cells were incubated at a density of 3.0×10^6^ cells/well for 7 days at 37 °C and 5% CO2 in ultra-low attachment six-well plates (Corning Life Sciences, Tewksbury, MA) to allow the differentiation of monocytes into macrophages. Then, the macrophages were verified to be approximately 95% CD68^+^ by flow cytometry. **(Figure S8).**

### Stimulation and polarization of primary human monocyte-derived macrophages

For qPCR and RNA sequencing analyses, human monocyte-derived macrophages were seeded in 24 well plates at a density of 6.5 x 10^5^ cells/well in R5 media. After overnight incubation at 37 °C and 5% CO2, the medium was replaced with fresh R5 media. The macrophages were either stimulated with 200 ng/ml soluble TRAIL (R&D, Minneapolis, MN) for 8 hours or pre-stimulated with 200 ng/ml TRAIL for 2 hours, and then in accordance with the literature polarized into M1 with 100 ng/mL LPS (Ultrapure; InvivoGen, San Diego, CA) and 20 ng/mL IFNγ (R&D, Minneapolis, MN); M2a with 20 ng/mL IL-4 (R&D, Minneapolis, MN) or M2c with 10 ng/mL IL-10 (R&D, Minneapolis, MN) for 6 hours (6, 67, 68). Control group macrophages were either left unstimulated (M0) or stimulated with the relevant polarization factors (M1, M2a, M2c) for 6 hours.

For the stimulation of DR4 (4C9) and DR5 (D269H/E195R) receptor-specific TRAIL mutants, M0 macrophages were treated with only DR4 mutant or only DR5 mutant, or both DR4 and DR5 mutants simultaneously for 8 hours. In M2 macrophage groups, cells were first pre-treated with only DR4 mutant or only DR5 mutant, or both DR4 and DR5 mutants for 2 hours, followed by polarization into M2a or M2c macrophages for 6 hours. Control group macrophages were either kept as unpolarized (M0) or polarized into M2a or M2c for 6 hours. These mutants were generated by protein engineering by altering the target receptor specificity of TRAIL (69, 70) and were obtained from Dr. Eva Szegezdi (University of Galway, Galway, Ireland).

For flow cytometry and ELISA analyses, the cells were stimulated in 48 well plates at a density of 3.0-3.5 x 10^5^ cells/well, in fresh R5 media. To assess the production of intracellular chemokines/cytokine by flow cytometry, macrophages were stimulated with TRAIL for 8 hours or pre-stimulated with TRAIL for 2 hours and then polarized into M1, M2a, and M2c macrophage subtypes for 6 hours while control group macrophages were either left unstimulated (M0) or stimulated with the related polarization factors (M1, M2a, M2c) for 6 hours. In the last 4 hours of polarization, all groups were treated with 0.5 µl/ml Brefeldin A (BD, Golgi Plug Protein Transport Inhibitor) and 0.33 µl/ml Monensin (BD, Golgi Stop Protein Transport Inhibitor). For the detection of remaining markers (cell surface markers and IDO) and viability of macrophages by flow cytometry, cells were either stimulated with TRAIL for 18 hours or pre-stimulated with TRAIL for 6 hours and then polarized with M1, M2a, or M2c polarization factors for 12 hours. Control group macrophages were either left unstimulated (M0) or stimulated with related polarization factors (M1, M2a, M2c) for 12 hours. Overall, in TRAIL or DR4/DR5 mutant-treated macrophage groups, TRAIL or DR4/DR5 mutants were retained in the culture media when the M1 or M2 polarization factors were introduced. The polarization factors remained in the media until the end of the incubation time. Both the control and TRAIL or DR4-DR5 mutant-treated macrophage groups were stimulated with M1 or M2 polarization factors for the same duration. To perform the mRNA level analyses, TRAIL or DR4/DR5-treated M1 and M2 macrophages were stimulated with TRAIL ligands for a total of 8 hours, including both the 2 hours of pre-treatment with TRAIL ligands and 6 hours of the polarization process. Similarly, for the protein-level analyses, TRAIL-treated M1 or M2 macrophages were incubated with TRAIL for a total of 18 hours (6 hours of pre-treatment and 12 hours of polarization). In TRAIL-treated M0 macrophages, TRAIL treatment was applied during the total time that TRAIL ligand is present in TRAIL-treated M1/M2 polarized macrophage groups. Therefore, M0 macrophages were stimulated with TRAIL or DR4/DR5 ligands for 8 hours or 18 hours, respectively.

For the assessment of the M2 to M1 switch by qPCR, first, macrophages were polarized into M2a and M2c for 2 hours and then stimulated with TRAIL for 6 hours. For the analysis at the protein level, macrophages were polarized first for 6 hours into M2a and M2c and then stimulated with TRAIL for 18 hours. Control group macrophages were either left unstimulated (M0) or polarized into M2a and M2c with the relevant factors for 8 or 24 hours, respectively. In this experimental setup, macrophages were treated with TRAIL after the polarization was initiated with M2 stimulants. The M2 polarization factors were retained in the culture media when TRAIL ligand was introduced. Both the control and TRAIL-treated groups were incubated with M2 stimulants for the same duration. Hence, control groups were treated with M2 stimulants for the total incubation time as the TRAIL-treated macrophage groups which are 8 hours (2 hours of initial polarization and 6 hours of TRAIL treatment) for the mRNA level analyses or 24 hours (6 hours of initial polarization and 18 hours of TRAIL treatment) for the protein level analyses.

### Tumor-associated macrophage (TAM) generation

U937 (AML), 5637 (bladder), and SKBR3 (breast) tumor cell lines were incubated in R10 culture media until the cell density reached 80-90 %. After reaching confluency, cells were washed twice with PBS and incubated for 24 h in fresh R10 culture media. Then, conditioned media (CM) of tumor cells was harvested by centrifugation at 400 g for 5 min. CM was filtered through a 0.22 µM filter (Millipore, MA, USA) to remove cell debris and stored at –20^0^C.

Tumor-associated macrophages (TAMs) were generated *in-vitro.* First, macrophages were seeded in 48 well plates at a density of 3.5 x 10^5^ cells/well and were incubated overnight. Then, cells were exposed to 1/2 diluted CM of tumor cell lines for 72 hours, and 200 ng/ml TRAIL ligand was introduced into the culture media in the last 18 hours of incubation without removing the tumor CM. Control groups were either left unstimulated or only stimulated with tumor CM for 72 hours.

### mRNA sequencing (RNA-seq)

According to the manufacturer’s recommendation, the RNA of macrophage groups was isolated with a Nucleo-Spin RNA kit (Macharey Nagel, Germany). The purity of RNA was checked with a NanoPhotometer® spectrophotometer (IMPLEN, CA, USA). RNA integrity and quantitation were assessed with Agilent 2100 Bioanalyzer system (Agilent Technologies, CA, USA) by using RNA 6000 Nano Assay Reagent Kit. RIN (RNA Integrity Number) scores of RNA samples were between 8.7 and 9.9.

RNA-Sequencing libraries were generated by using NEBNext® UltraTM RNA Library Prep Kit for Illumina® (NEB, USA) according to the manufacturer’s recommendations and index codes were added to each sample. First, the positive selection of polyA+ RNA with poly-T oligo-attached magnetic beads and the removal of rRNA with the Ribo-Zero kit was performed. Then, the library construction was followed by the fragmentation of mRNA, synthesis of the first and second strands of cDNA, ligation of the adaptors, and performing an enrichment process. After the clustering of the index-coded samples with cBot Cluster Generation System by using TruSeq PE Cluster Kit v3-cBot-HS (Illumia), library sequencing was performed by ILLUMINA NextSeq 500 instrument as 40 million 150-bp paired-end reads per sample.

Quality control of the samples, preparation of libraries, and sequencing were performed by Novogene (UK) Company Limited.

### RNA-seq data analysis

The quality control of reads from sequencing libraries was performed with FASTQC tool (v0.11.9; https://www.bioinformatics.babraham.ac.uk/projects/fastqc/). Adapter removal and quality trimming of reads were done with Trimmomatic (v0.39) (71). The Human reference genome (GRCh38) in FASTA format and associated gene annotation in General Transfer Format (GTF) were obtained from the Ensembl website (Release 99; https://www.ensembl.org/). Sequencing libraries were aligned to the human reference genome with Rusbread v2.0.3 package (72) of R v3.5.1 statistical computing environment (https://www.r-project.org/) with the following settings: *“align(index={index file}, readfile1={input_1.fastq}, readfile2={input 2.fastq} type=“rna”, input_format=“gzFASTQ”, output_format=“BAM”, output_file={output file}, nthreads=numParallelJobs)”*. SAMtools v1.3.1 (73) was utilized to sort and index the BAM files generated in the alignment step. To measure the expression levels of genomic features, the featureCounts function (74) of Rsubread package with the following command: “featureCounts (files = {infile.bam}, annot.ext = “{infile.gtf}”, isGTFAnnotationFile = T, GTF.featureType = “exon”, GTF.attrType = “gene_id”, useMetaFeatures = T, countMultiMappingReads = T, isPairedEnd = T, nthreads = numParallelJobs)” was employed. Fragments Per Kilobase Million (FPKM) values of each feature across samples were calculated with “rpkm” function of edgeR package (v3.24.3) (75). We removed the features where expression levels are < 1 FPKM in each group, and only considered samples where at least one group has FPKM **≥** 1 threshold in at least three out of four donors. To identify the genomics features differentially expressed between the groups, we again utilized edgeR package. In this analysis step, Trimmed Mean of M-values (TMM) normalization was employed to the filtered count values and the dispersions were estimated with estimateDisp function for each pairwise comparison. To calculate False Discovery Rate (FDR) value of each feature, proper contrast statistics were employed with “exactTest” function of the package. The genomic features were considered as differentially expressed when the absolute value of log2-fold change was **≥** 0.6 (equal or greater than ∼1.5 fold change) and FDR-value < 0.05.

### Statistical analysis of RNA seq and generation of pathway and PCA plots

R statistical computation environment was utilized for all statistical analyses. ClusterProfiler v3.18.0 (76) was used to explore and visualize the enrichment of Kyoto Encyclopedia of Genes and Genomes (KEGG) terms in the sets of genes of interest. The top 10 significantly enriched pathways of differentially expressed genes between TRAIL-treated vs control samples were depicted. The R program package clusterProfiler (76, 77) was used to perform KEGG pathway analysis and functional annotation for differentially expressed genes. 238, 157 and 164 KEGG pathways were mapped for M0, M2a and M2c groups, respectively (p <0.05).

We also employed pheatmap package of R (https://CRAN.R-project.org/package=pheatmap/) to draw all the heatmaps, on which expression values were represented in the rows. Clustering was performed with the Euclidian method and prcomp function was used for the Principal Component Analysis (PCA). Other graphics were obtained using the ggplot2 package (https://ggplot2.tidyverse.org/).

### Ovarian and sarcoma cancer dataset analysis

#### Survival analysis

Kaplan-Meier plots were generated to examine the correlation between TRAIL (TNFSF10) gene expression and overall survival of ovarian and sarcoma cancer patients using the TCGA database with the KM plotter tool (78). Gene expression data and corresponding clinical data for 373 ovarian cancer and 259 sarcoma cancer patients were extracted. The cases were divided into high and low macrophage content by using xCell algorithm (79). The median expression of TRAIL across all patients within each tumor type was used to categorize patients into high and low TRAIL expression groups. Log rank test was applied to calculate hazard ratio (HR), p-values, and 95% confidence intervals (CI) for each group.

#### Correlation analysis

TCGA datasets of ovarian and sarcoma cancer transcriptomes were used to extract gene expression data and generate the correlation plots by using cBioPortal (80, 81). Spearman’s rank and Pearson correlation analyses were used to examine the relationship between TRAIL gene expression and M1 marker gene (CXCL10, CXCL11, IFI44L, and CD38) expression or M2 marker gene (TGM2, HGF, HPGD, and FAXDC2) expression. The statistical significance level was chosen as p < 0.05.

#### Proteomics analysis

Clinical Proteomic Tumor Analysis Consortium (https://cptac-data-portal.georgetown.edu/cptacPublic/) database was used to obtain the processed protein expression data of ovarian cancer patients with the accession number PDC000113. The selected proteins were filtered, and R statistical computation environment was utilized to perform statistical analysis and generate correlation plots. Spearman’s rank and Pearson correlation analyses were used to demonstrate the association between TRAIL expression and M1 marker (CD38 and IFI44L) expression or M2 marker (TGM2 and HPGD) expression at the protein level. A p-value < 0.05 was considered statistically significant.

### Quantitative PCR (qPCR)

The total RNA of macrophage groups was isolated with a Nucleo-Spin RNA kit (Macharey Nagel, Germany) or Monarch Total RNA Miniprep Kit (New England Biolabs, Massachusetts, ABD). The purity and quantity of RNA were evaluated by a Nanodrop spectrophotometer (ThermoFisher Scientific, Waltham, MA, USA). The amount of RNA was adjusted to a minimum of 600 ng in each sample and cDNA synthesis was performed with EvoScript Universal cDNA Master (Roche, Basel, Switzerland) according to the manufacturer’s protocol.

For the verification of RNA sequencing analysis, Fast Start Essential DNA Green Master (Roche, Basel, Switzerland) and Sybr Green QuantiTect Primer Assays (Qiagen, Hilden, Germany) were used for CXCL10 (#QT01003065), CXCL1 (#QT00199752, CXCL11 (#QT02394644), CCL15 (#QT00032165), IL1B (#QT00021385), CD38 (#QT00073192), IDO1 (#QT00000504), ACOD1 (#QT01530424), IFI44L (#QT00051457), IL12B (#QT00000364); TMEM37 (QT01530361), FAXDC2 (#QT01678348), HTR2B (#QT00060368), MS4A6A (QT00097377), HPGD (#QT00013454), ANGPTL4 (#QT00003661), SELENOP (#QT01008175), HGF (#QT00065695), PRR5L (#QT00200998), F13A1 (#QT00042700), and ACTB (#QT00095431). For the detection of classical M1 and M2 markers, Fast Start Essential DNA Probes Master (Roche, Basel, Switzerland) and RealTime ready Single Assays (Roche, Basel, Switzerland) were used for CXCL10 (#100134759), TNF (#100134777), IDO1 (#100134768), CXCL9 (#100137998); MRC1 (#100134731), TGM2 (#100134722), CD23 (#100125140), CCL17 (#100138007), CCL22 (#100134713), IL-10 (#100133437), CD163 (#100134801), and ACTB (#100074091).

LightCycler 480 II Real-Time System (Roche, Basel, Switzerland), or Applied Biosystems 7500 Fast Real-Time PCR System (ThermoFisherScientific, Waltham, MA, USA) was used to perform qPCR analyses. Each step was carried out as recommended by the manufacturers. ACTB was used as the reference gene. The relative quantification of gene expression was determined by the 2^-ΔΔCT^ method (82).

## ELISA

Supernatants of samples were collected and stored at –20^0^C until further usage. The concentrations of M1 cytokines/chemokine [TNF (Invitrogen, ThermoFisher Scientific, Waltham, MA, USA), IL-12 p70 (BioLegend, San Diego, CA), IL-1β (BioLegend, San Diego, CA), CXCL11 (R&D, Minneapolis, MN)], and M2 cytokines [(IL-10 (BioLegend, San Diego, CA), HGF (R&D, Minneapolis, MN)] were determined by ELISA in accordance with the manufacturer’s instructions. The absorbance of each sample was measured at 450 nm and 570 nm wavelengths.

### Flow cytometry

After collecting the supernatants of samples and washing them with PBS once, StemPro Accutase Cell Dissociation Reagent (Gibco; ThermoFisher Scientific, Waltham, MA, USA) was used to detach macrophages according to the manufacturer’s instructions. Zombie UV Fixable Viability Kit (BioLegend, San Diego, CA, USA) was used to detect and eliminate dead cells. The cells were incubated in flow cytometry staining buffer (PBS, 1% bovine serum albumin, 0.1% sodium azide) containing Human TruStain FcX antibody (Fc block) (BioLegend, San Diego, CA) for 15 minutes on ice. Subsequently, cells were stained with anti-human antibodies as indicated below.

For surface staining, cells were incubated in flow cytometry staining (FACS) buffer containing; CD86-BV605 (IT2.2; 1:200), HLA-DR-APC-Cy7 (L243; 1:200), HLA-DR-PE (L243; 1:400), CD64-PerCP-Cy5.5 (10.1; 1:200), CD200R-PE-Dazzle594 (OX-108; 1:200), CD206-AF700 (15-2; 1:200), CD206-BV421 (15-2; 1:100), CD163-PE-Cy7 (GHI/61; 1:200), CD38-BV510 (HB-7; 1:100), DR4-APC (DJR1; 1:100), DR5-PE (DJR2-4; 1:100) antibodies for 45 minutes on ice in the dark. Antibodies were purchased from BioLegend (San Diego, CA, USA). For intracellular staining, cells were fixed with Cytofix-Cytoperm (BD Biosciences, USA) solution for 15 minutes on ice. Then, cells were washed with Perm/Wash solution (BD Biosciences, USA), and then incubated in the same buffer containing CXCL10-AF-647 (33036; 1:50), CXCL1-AF-405 (20326; 1:40), IDO-PE (700838; 1:25), TNF-PE (MAb11; 1:80), CD68-FITC (Y1/82A; 1:100) antibodies for 45 minutes at room temperature in the dark. CXCL10, CXCL1, and IDO antibodies were purchased from R&D (Minneapolis, MN) and TNF antibody was purchased from e-Biosciences (Santa Clara, CA, USA). CD68 antibody was purchased from Biolegend (San Diego, CA, USA). After the staining, samples were washed, suspended in FACS buffer, and analyzed by flow cytometry.

For the analysis of macrophage viability, after the detachment of cells, samples were washed twice with FACS buffer. Then, cells were incubated in Annexin V Binding Buffer containing Annexin V-PE and 7-AAD (640934; 1:20) according to the manufacturer’s instructions (BioLegend, San Diego, CA) for 15-20 minutes at room temperature. After the incubation staining was terminated, and cells were directly analyzed by flow cytometry.

In all protocols, cells were stained in V-bottom 96-well plates (Greiner Bio-One, North Carolina, USA), and before assessment by flow cytometry, they were transferred in FACS tubes (Greiner Bio-One, North Carolina, USA). Fluorescence emissions were detected by LSR Fortessa (BD Biosciences) and the data were analyzed with the FlowJo software (TreeStar, Ashland, OR).

### Co-culture of macrophages and AML tumor cells

U937 AML tumor cell line was used to analyze macrophage cytotoxicity. AML cells were cultured in R10 media [(RPMI 1640 supplemented with 10% heat-inactivated fetal bovine serum, 1% L-glutamine, 1% Penicillin-Streptomycin, 1% Non-essential amino acids (Gibco, ThermoFisher Scientific, Waltham, MA, USA) and 1% Na-pyruvate (Gibco, ThermoFisher Scientific, Waltham, MA, USA)] at a density of 3.0 x 10^5^ cells/10 ml in T25 cell culture flasks (Sarstedt, Numbrecht, Germany). After reaching confluency, cells were passaged every 2 days.

For the co-culture, first labeled primary human macrophages were seeded in 48 well plates at a density of 3.5 x 10^5^ cells/well and were incubated overnight. Then, cells were treated with 200 ng/ml TRAIL for 6 hours and polarized into M1 with 100 ng/mL LPS and 20 ng/mL IFNγ for 12 hours in fresh R5 media or were only stimulated with M1 stimulants for 12 hours. After that, the cells were washed once with fresh media. At this stage, labeled AML cells were seeded on the macrophages in fresh R10 media at a 10:1 effector (macrophages) to target (AML) cell ratio, and cells were incubated for 72 hours. Control AML groups were cultured in the absence of macrophages in R10 media for 72 hours.

Macrophages were labeled with 1μM CFSE (BioLegend, San Diego, CA), and for the staining of AML cells, 2.5 μM Tag it-Violet (BioLegend, San Diego, CA) was used. Stainings were performed by the manufacturer’s instructions.

### Macrophage cytotoxicity and phagocytosis assays

At the end of the co-culture, supernatants were collected into FACS tubes (Greiner Bio-One, North Carolina, USA), and cells were washed with PBS once. As the manufacturer suggested, TrypLE cell dissociation reagent (Gibco, ThermoFisher Scientific, Waltham, MA, USA) was used to detach cells. Then, cells were collected, transferred into FACS tubes, and washed twice with PBS. At the last step, they were stained with 5 nM SYTOX Red in PBS containing 0.5% BSA (ThermoFisher Scientific, Waltham, MA, USA) for 15 minutes on ice in the dark. Subsequently, both the cytotoxicity of macrophages for AML cells (Tag-it Violet^+^ CFSE^-^ Sytox Red^+^) and the phagocytosis of AML cells by live macrophages (CFSE^+^ Sytox Red^-^ Tag-it Violet^+^) were assessed by Canto II (BD Biosciences). The data were analyzed with the FlowJo software (TreeStar, Ashland, OR). Macrophage cytotoxicity for tumor cells was calculated as % of Sytox Red^+^ AML cells (Tag-it Violet^+^ CFSE^-^) co-cultured with macrophages (CFSE^+^)-% Sytox Red^+^ AML cells alone.

### Statistical Analysis

Graph Pad Prism 8 (GraphPad Software, CA, USA) was used to analyze data and generate graphs. First, the normal distribution of the data was analyzed with the Shapiro-Wilk test to perform biostatistical analyses. Then, the calculation of p-values was performed by a two-tailed paired Student’s t-test, Wilcoxon matched-pairs signed-rank test; One-way ANOVA test, or Friedman test according to the experimental groups and the normal data distribution. For the normal data distribution, Student’s t-test, or One-way ANOVA test, was applied and the results were shown as mean ± SEM. For the non-normal data distribution, Wilcoxon matched-pairs or Friedman test, were applied and the results were shown as median with interquartile range. Data containing two different groups were analyzed with Student’s t-test or Wilcoxon matched-pairs signed-rank. If data contains more than two groups One-Way Anova test or Friedman test was used. P values less than 0.05 were considered statistically significant.

## RESULTS

### The impact of TRAIL on the viability of primary human macrophage subtypes

TRAIL has selective cytotoxicity on cancer cells but it can also induce cell death in M1 or M2 macrophage subtypes according to the environment (34, 47, 48, 60, 83, 84). Therefore, first, the impact of soluble TRAIL on the viability of unpolarized (M0), and M1 and M2 polarized primary human macrophages was investigated. TRAIL did not induce cell death in primary human M1 and M2c macrophage subtypes **(Figure 1B).** Regarding M0 and M2a macrophages, it slightly decreased (6,43% and 4,34% respectively) the percentage of live cells (Annexin V-/7AAD-) **(Figure 1B).** These data show that TRAIL does not majorly affect the viability of M0 and M1/M2 polarized macrophages.

**Figure 1.**
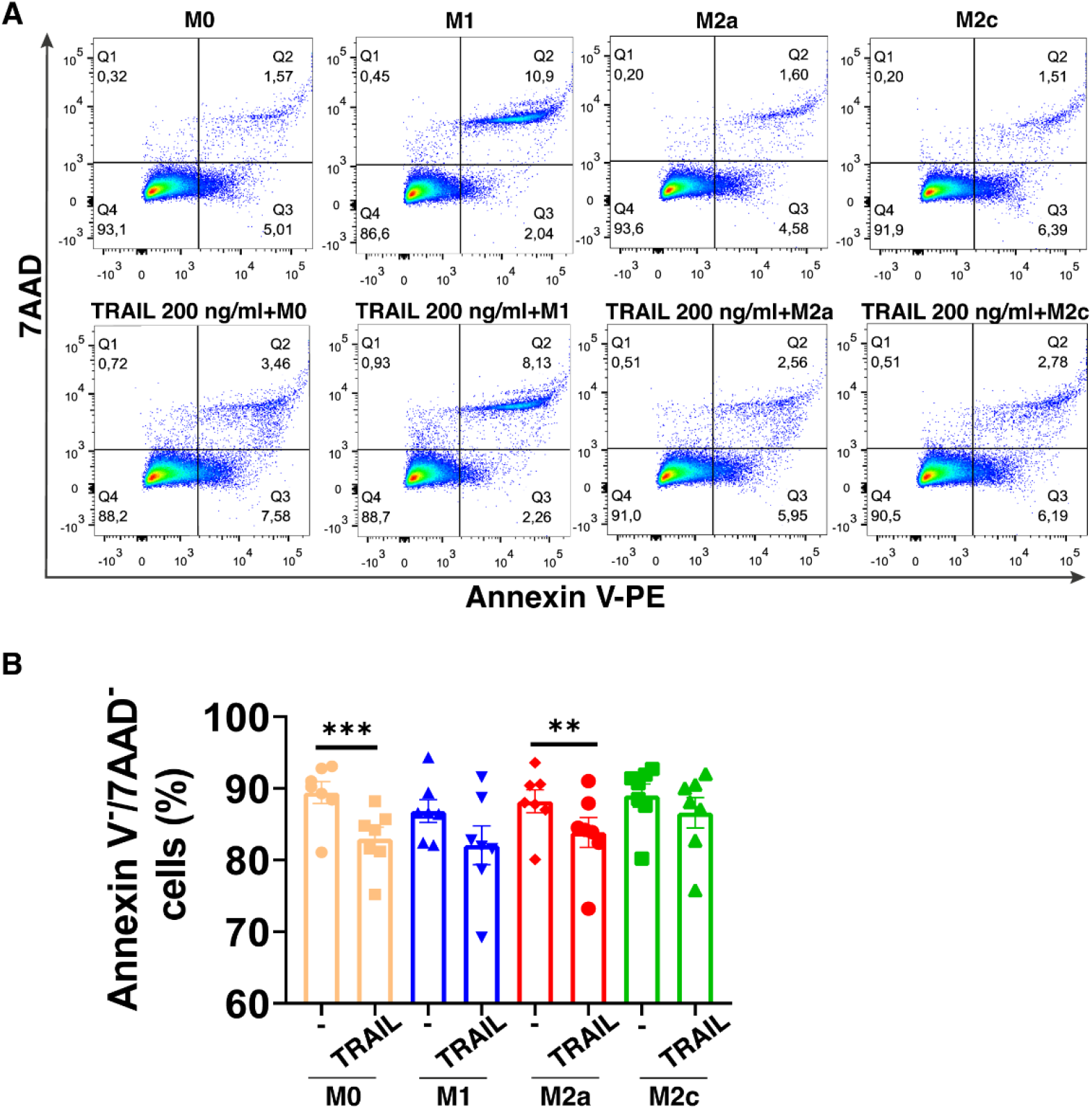
TRAIL does not majorly affect the viability of primary human macrophage subtypes. M0 macrophages were stimulated with 200 ng/ml TRAIL for 18 hours. For polarization, macrophages were pre-stimulated with TRAIL for 6 hours and then polarized into M1 (100 ng/ml LPS and 20 ng/ml IFNγ), M2a (20 ng/ml IL-4), or M2c (10 ng/ml IL-10) for 12 hours. Control groups were left unstimulated or stimulated with only M1, M2a, or M2c polarization factors for 12 hours. Cell viability was analyzed with Annexin V/ 7-AAD staining by flow cytometry. (A) Representative dot plots and (B) Bar graphs show the percentage of live macrophages (AnnexinV^-^7AAD^-^). Data shown are mean ± SEM pooled from two independent experiments (n=7). Statistical analyses were performed with a One-way ANOVA with Sidak’s multiple comparisons post-hoc test between untreated and TRAIL-treated macrophages, polarized and TRAIL-treated polarized macrophages, **P<0.01 and ***P<0.001.

### The impact of TRAIL on the transcriptome of primary human macrophage subtypes

RNA sequencing analysis was applied to perform an in-depth analysis of the impact of TRAIL on the polarization of primary human macrophages. Differential expression of M1/M2 markers between TRAIL-treated M0, M1, M2a, and M2c macrophages and control groups was compared.

Principal component analysis (PCA) demonstrated that independent of TRAIL treatment, M1 macrophages clustered separately from the other macrophage subtypes while M2a macrophages showed a slight transcriptomic signature difference from M0 and M2c macrophages **(Figure 2A)**.

**Figure 2.**
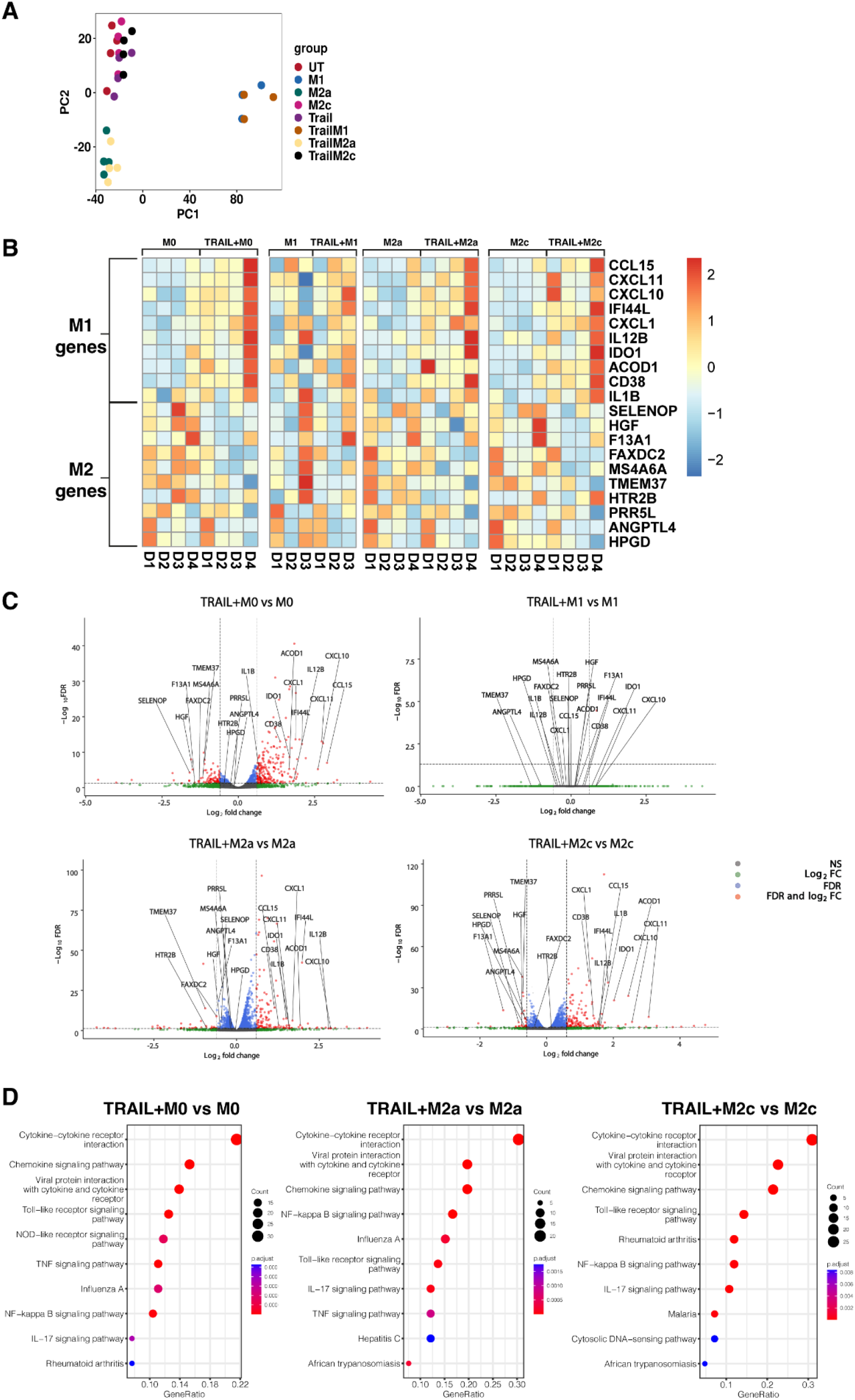
TRAIL increases the differential expression of M1 markers while decreasing the expression of M2 markers in primary human M0, M2a, and M2c macrophage subtypes. M0 macrophages were stimulated with 200 ng/ml TRAIL for 8 hours. For polarization, macrophages were pre-stimulated with TRAIL for 2 hours and then polarized into M1 (100 ng/ml LPS and 20 ng/ml IFNγ), M2a (20 ng/ml IL-4), or M2c (10 ng/ml IL-10) for 6 hours. Control groups were left unstimulated (UT) or stimulated with only M1, M2a, or M2c polarization factors for 6 hours (n=3-4). Differential gene expression was analyzed by RNA sequencing. (A) PCA plot of transcriptomic profiles of TRAIL-treated and control macrophage subtypes were shown. Selected M1 and M2 markers in each macrophage subtype were shown as (B) heat maps and (C) volcano plots comparing TRAIL-treated and control macrophages. (D) KEGG analysis on differentially expressed genes (FDR value ≤0.05, log2FC value ≥ 0.6/ FC ≥ 1.5) comparing TRAIL-treated and control macrophages was shown and the top 10 significant pathways were indicated for each subtype. Data were z-score normalized for heatmaps. D: Donor

Even though TRAIL-treated and control group macrophages stand in the same cluster, TRAIL-induced the differential expression (FDR value ≤0.05, log2FC value ≥ 0.6/ FC ≥ 1.5) of certain M1 and M2 markers in macrophage subtypes **(Figure 2B**, **Figure 2C).**

As demonstrated in heatmaps and volcano plots, the expression of the classical M1 markers CXCL1 (85), CXCL11, CXCL10, IDO1, IL1B, IL12B (17) and CCL15 (86), and the more recently accepted M1 markers CD38 (87), ACOD1 (88) and IFI44L (89) were induced by TRAIL in M0, M2a, and M2c macrophages. All of these markers are known to be upregulated by inflammatory agents such as LPS and IFNγ in primary human/mouse macrophages and THP-1 human macrophage cell line (87, 90, 91, 92, 93, 94, 95, 96, 97, 98, 99, 100, 101, 102, 103). Furthermore, our results demonstrated that the expression of the recently identified M2 markers F13A1, MS4A6A (95), TMEM37 (104), FAXDC2 (105), SELENOP (106, 107), HGF (108), PRR5L (109), HPGD (110), ANGPTL4 (111), and HTR2B (112) were decreased by TRAIL in M0, M2a, and M2c macrophages **(Figure 2B, 2C, and Table S1A, S1C, S1D)**. All of these M2 markers are known to be induced by regulatory mediators such as IL-4, IL-10, or PPARγ ligands, while downregulated in the presence of inflammatory factors in primary human and mouse macrophages (5, 105, 106, 108, 109, 113, 114, 115, 116, 117, 118, 119, 120, 121). However, in M1 macrophages, TRAIL did not induce a significant change in the expression of these M1 or M2 markers **(Figure 2B, 2C, Table S1B).**

Next, the Kyoto Encyclopedia of Genes and Genomes (KEGG) analysis was performed to characterize the molecular pathways in M0, M2a, and M2c macrophages after TRAIL treatment. In all macrophage subtypes, several immune-related pathways were upregulated with TRAIL, and among them, cytokine-cytokine receptor interaction was identified as the most enriched. Besides, TRAIL stimulation was associated with inflammatory pathways such as NFkB, IL-17, TLR, and TNF signaling pathways in respective macrophage subtypes. **(Figure 2D).**

Next, the differential expression of M1 and M2 markers in M0, M2a, and M2c macrophage subtypes were verified by qPCR. Consistent with the RNA sequencing analysis, TRAIL treatment increased the expression of all the M1 markers except IL12B, while decreasing the expression of all the M2 markers except ANGPTL4 and F13A1 in primary human macrophage subtypes at the mRNA level **(Figure 3).** These results show that TRAIL effectively impacts the expression of both M1 and M2 markers in macrophage subtypes at the transcriptomic level by promoting the induction of M1 markers while downregulating the expression of M2 markers.

**Figure 3.**
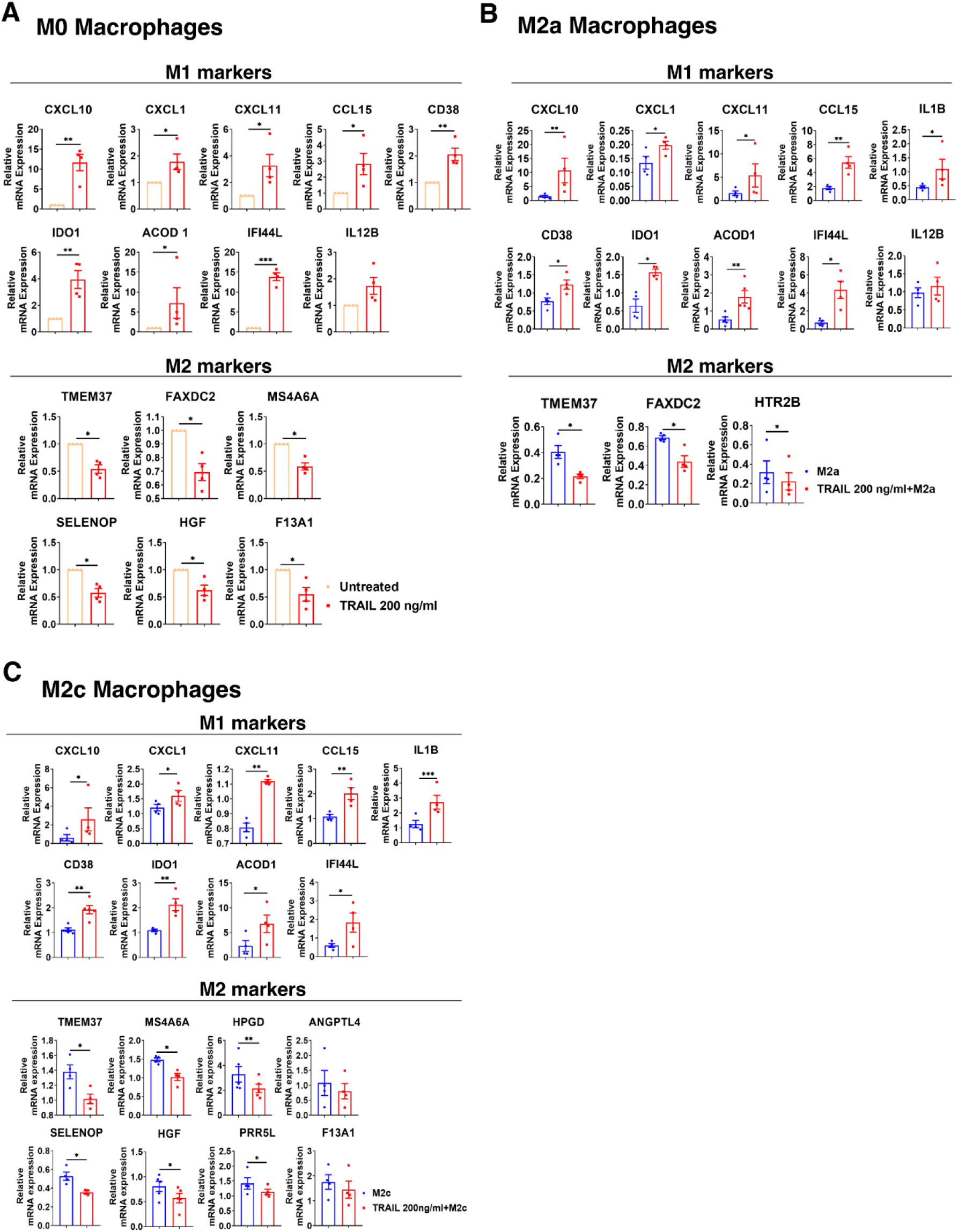
TRAIL increases the expression of M1 markers while decreasing the expression of M2 markers in primary human M0, M2a, and M2c macrophage subtypes at the mRNA level. M0 macrophages were stimulated with 200 ng/ml TRAIL for 8 hours. For polarization, macrophages were pre-stimulated with TRAIL for 2 hours and then polarized into M1 (100 ng/ml LPS and 20 ng/ml IFNγ), M2a (20 ng/ml IL-4), or M2c (10 ng/ml IL-10) for 6 hours. Control groups were left unstimulated or stimulated with only M1, M2a, or M2c polarization factors for 6 hours. qPCR analyses of M1 and M2 markers in (A) M0, (B) M2a, and (C) M2c macrophage subtypes were shown. Data shown are mean ± SEM pooled from two independent experiments (n=4-5). Statistical analyses were performed with a two-tailed paired Student’s t-test between control and TRAIL-treated groups, *P<0.05, **P<0.01, ***P<0.001.

### The translational output of verified TRAIL-induced M1 markers in primary human macrophage subtypes

Among the verified M1 markers, CXCL10, CXCL1, IDO1, CD38, and CXCL11 were selected and analyzed at the protein level in TRAIL-treated M0, M2a, and M2c macrophage subtypes. The effect of TRAIL was also reflected at the protein level. Consistent with the changes at the mRNA level, TRAIL stimulation increased the production of most of the indicated M1 markers in primary human M0 and M2 macrophages at the protein level **(Figure 4-6)**. Therefore, our data show that TRAIL promotes M1 response by increasing the expression of M1 markers in unpolarized (M0) and M2 polarized (M2a, M2c) primary human macrophages both at mRNA and protein levels **(Figure 3**, **Figure 4-6)**.

**Figure 4.**
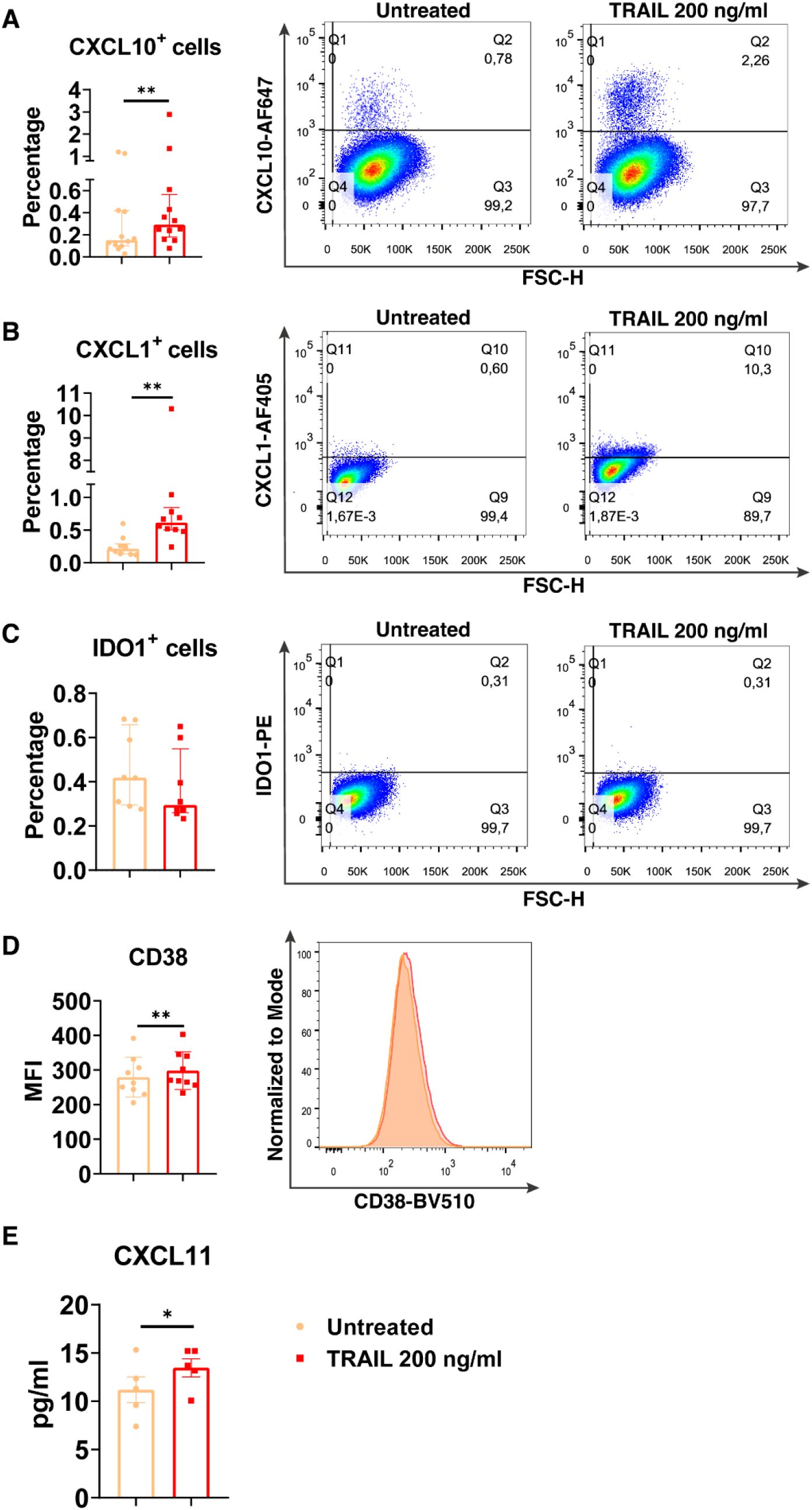
TRAIL increases the expression of M1 markers in primary human M0 macrophages at the protein level. M0 macrophages were stimulated with 200 ng/ml TRAIL for 8 hours (CXCL10/CXCL1) or 18 hours (IDO/CD38/CXCL11). Control groups were left unstimulated. (A-D) The production of CXCL10, CXCL1, IDO, and CD38 was analyzed by flow cytometry, and the representative plots are included. (E) The production of CXCL11 was analyzed by ELISA. Data shown are mean ± SEM or median with interquartile range pooled from two or more independent experiments [(A) n=11, (B) n=10, (C) n=8, (D) n=9, (E) n= 5]. Statistical analyses were performed with a two-tailed paired Student’s t-test or Wilcoxon matched-pairs signed-rank test between untreated and TRAIL-treated macrophages, *P<0.05, **P<0.01.

**Figure 5.**
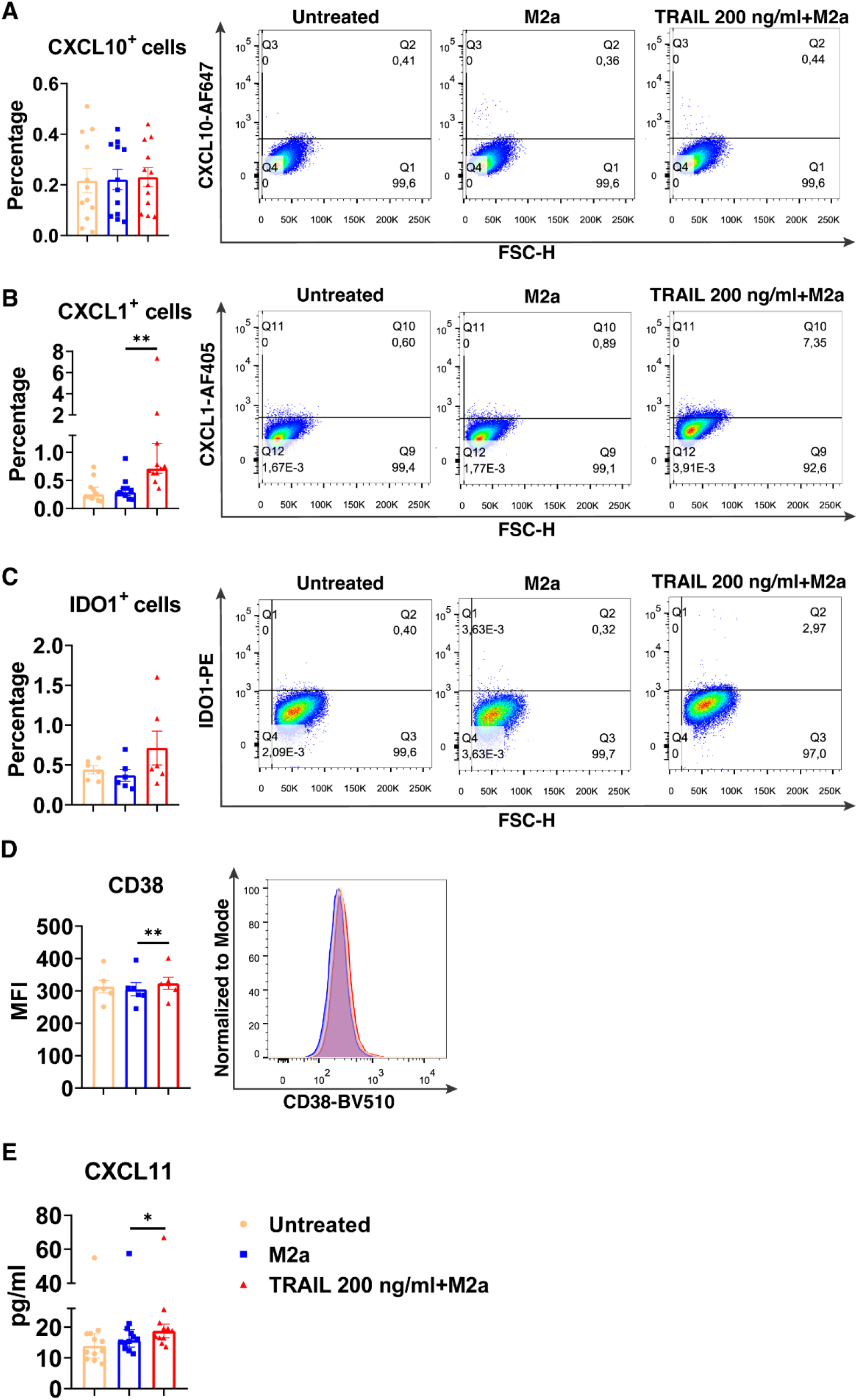
TRAIL increases the expression of M1 markers in primary human M2a macrophages at the protein level. Macrophages were pre-stimulated with 200 ng/ml TRAIL for 2 hours and then polarized into M2a with 20 ng/ml IL-4 for 6 hours for the analyses of the CXCL10 and CXCL1. For the remaining markers, macrophages were pre-stimulated with TRAIL for 6 hours and then polarized with IL-4 for 12 hours. Control groups were left unstimulated or stimulated only with 20 ng/ml IL-4 for 6 or 12 hours, respectively. (A-D) The production of CXCL10, CXCL1, IDO, and CD38 was analyzed by flow cytometry, and the representative plots are included. (E) The production of CXCL11 was analyzed by ELISA. Data shown are mean ± SEM or median with interquartile range pooled from two or more independent experiments [(A) n=12, (B) n= 11, (C) n=6, (D) n=6, (E) n= 12]. Statistical analyses were performed with a One-way ANOVA with Sidak’s multiple comparisons post-hoc test, or Friedman with Dunn’s multiple comparisons post-hoc test between untreated and M2a macrophages, M2a and TRAIL-treated M2a macrophages, *P<0.05, **P<0.01, ***P<0.001.

**Figure 6.**
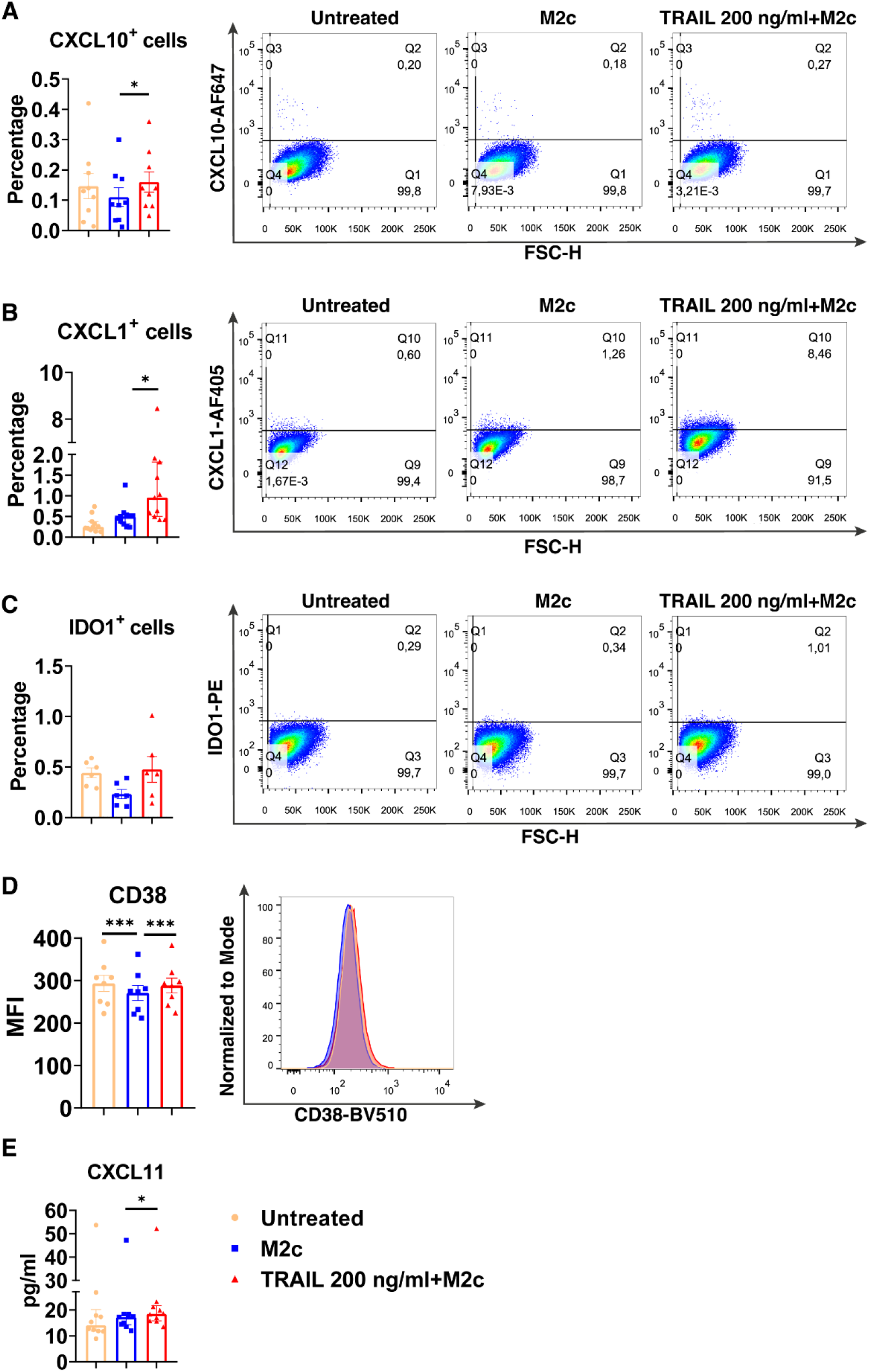
TRAIL increases the expression of M1 markers in primary human M2c macrophages at the protein level. Macrophages were pre-stimulated with 200 ng/ml TRAIL for 2 hours and then polarized into M2c with 10 ng/ml IL-10 for 6 hours for the analyses of the CXCL10 and CXCL1. For the remaining markers, Macrophages were pre-stimulated with TRAIL for 6 hours and then polarized with IL-10 for 12 hours. Control groups were left unstimulated or stimulated only with 10 ng/ml IL-10 for 6 or 12 hours, respectively. (A-D) The production of CXCL10, CXCL1, IDO, and CD38 was analyzed by flow cytometry, and the representative plots are included. (E) The production of CXCL11 was analyzed by ELISA. Data shown are mean ± SEM or median with interquartile range pooled from two or more independent experiments [(A) n=9, (B) n=11, (C) n=6, (D) n=8, (E) n= 10]. Statistical analyses were performed with a One-way ANOVA with Sidak’s multiple comparisons post-hoc test, or Friedman with Dunn’s multiple comparisons post-hoc test between untreated and M2c macrophages, M2c and TRAIL-treated M2c macrophages, *P<0.05, **P<0.01, ***P<0.001.

### The impact of TRAIL on the expression of classical M1 and M2 markers in primary human macrophage subtypes

After establishing the TRAIL-mediated differentially expressed markers in human macrophage subtypes, the expression of classical M1 and M2 markers, which are widely used in the literature (7, 95, 122, 123, 124, 125), was also analyzed.

Intriguingly, TRAIL increased the expression of classical M1 markers CXCL10, TNF, IDO1, and CXCL9 in M1 macrophages at the mRNA level **(Figure 7A)**. Furthermore, it increased the production of classical M1 cell surface activation markers CD86, HLA-DR alpha, and CD64; chemokines CXCL10 and CXCL1; and cytokines TNF, IL-1β, and IL-12-p70 in M1 macrophages at protein level **(Figure 7B)**. Besides, the percentage of M1 chemokine producing cell populations was increased in M1 macrophages upon TRAIL treatment **(Figure S3A and Figure S3C).** These data show that, even though RNA-seq analysis did not show a significant change in M1 markers in M1 macrophages after TRAIL treatment, TRAIL stimulation increases the expression of classical M1 markers both at mRNA and protein levels in this macrophage subtype **(Figure 7)**.

**Figure 7.**
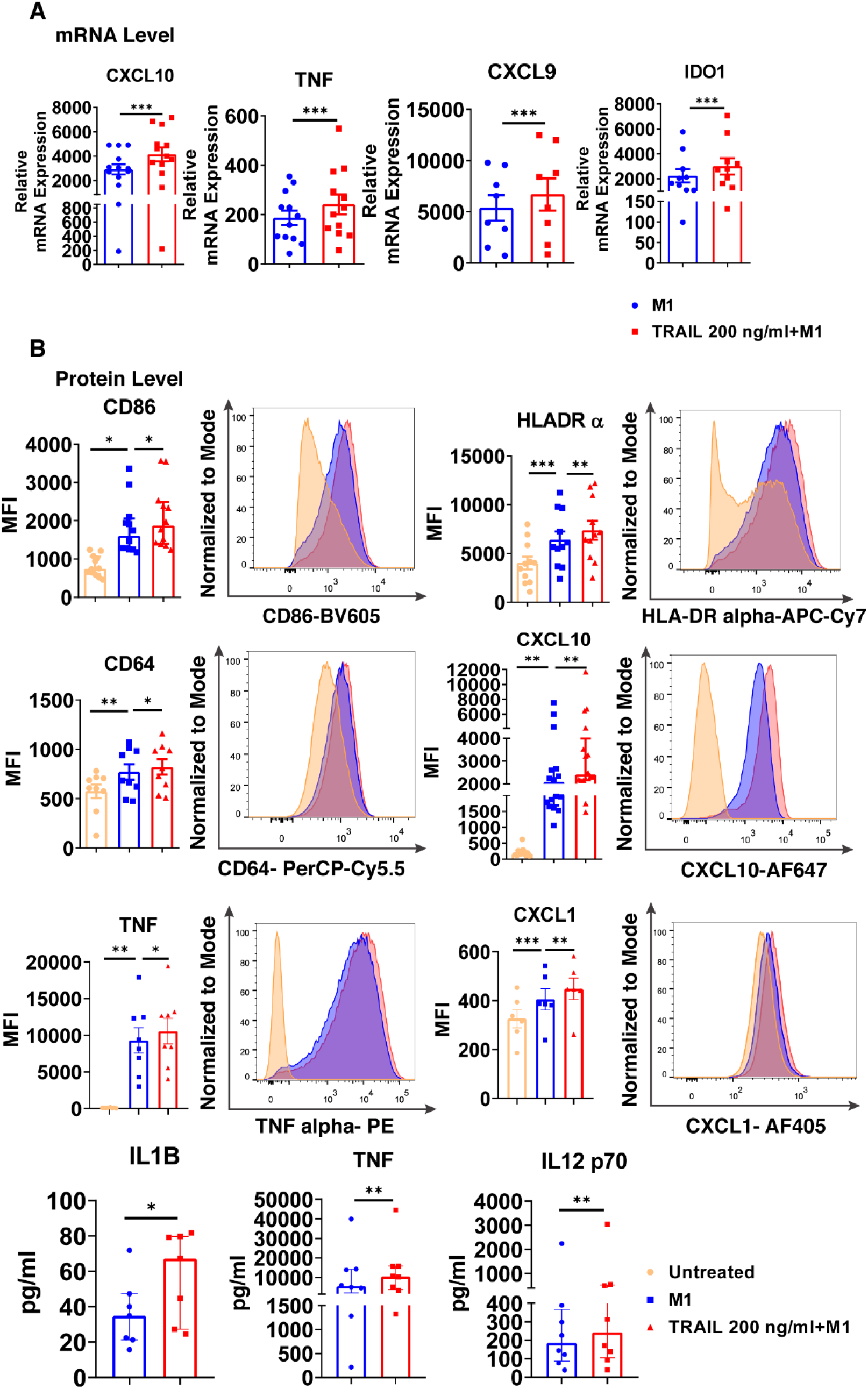
TRAIL increases the expression of classical M1 markers in primary human M1 macrophages at mRNA and protein levels. (A) Macrophages were pre-stimulated with 200 ng/ml TRAIL for 2 hours and then polarized into M1 with 100 ng/ml LPS and 20 ng/ml IFNγ for 6 hours. Control group was stimulated only with LPS and IFNγ for 6 hours. Expression of M1 markers was analyzed by qPCR. (B) Macrophages were pre-stimulated with 200 ng/ml TRAIL for 6 hours and then polarized into M1 with 100 ng/ml LPS and 20 ng/ml IFNγ for 12 hours. Control groups were left unstimulated or stimulated only with LPS and IFNγ for 12 hours. Expression of CD86, HLA-DR alpha, CD64, CXCL10, TNF, and CXCL1 was analyzed by flow cytometry, and representative plots are included. Production of IL-1β, TNF, and IL-12 p70 was analyzed by ELISA. Data shown are mean ± SEM or median with interquartile range pooled from two or more independent experiments [(A) n=8-13, (B) n= 6-16]. Statistical analyses were performed with a two-tailed paired Student’s t-test, Wilcoxon matched-pairs signed-rank test, One-way ANOVA with Sidak’s multiple comparisons post-hoc test, or Friedman with Dunn’s multiple comparisons post-hoc test between untreated and M1 macrophages, M1 and TRAIL-treated M1 macrophages, *P<0.05, **P<0.01, ***P<0.001.

Regarding M2a macrophages, TRAIL decreased the expression of the M2 markers MRC1, CD23, and TGM2, while increasing the chemokines CCL22 and CCL17 at the mRNA level **(Figure 8A)**. On the other hand, TRAIL did not affect the expression of M2a markers at the protein level **(Figure 8B)**.

**Figure 8.**
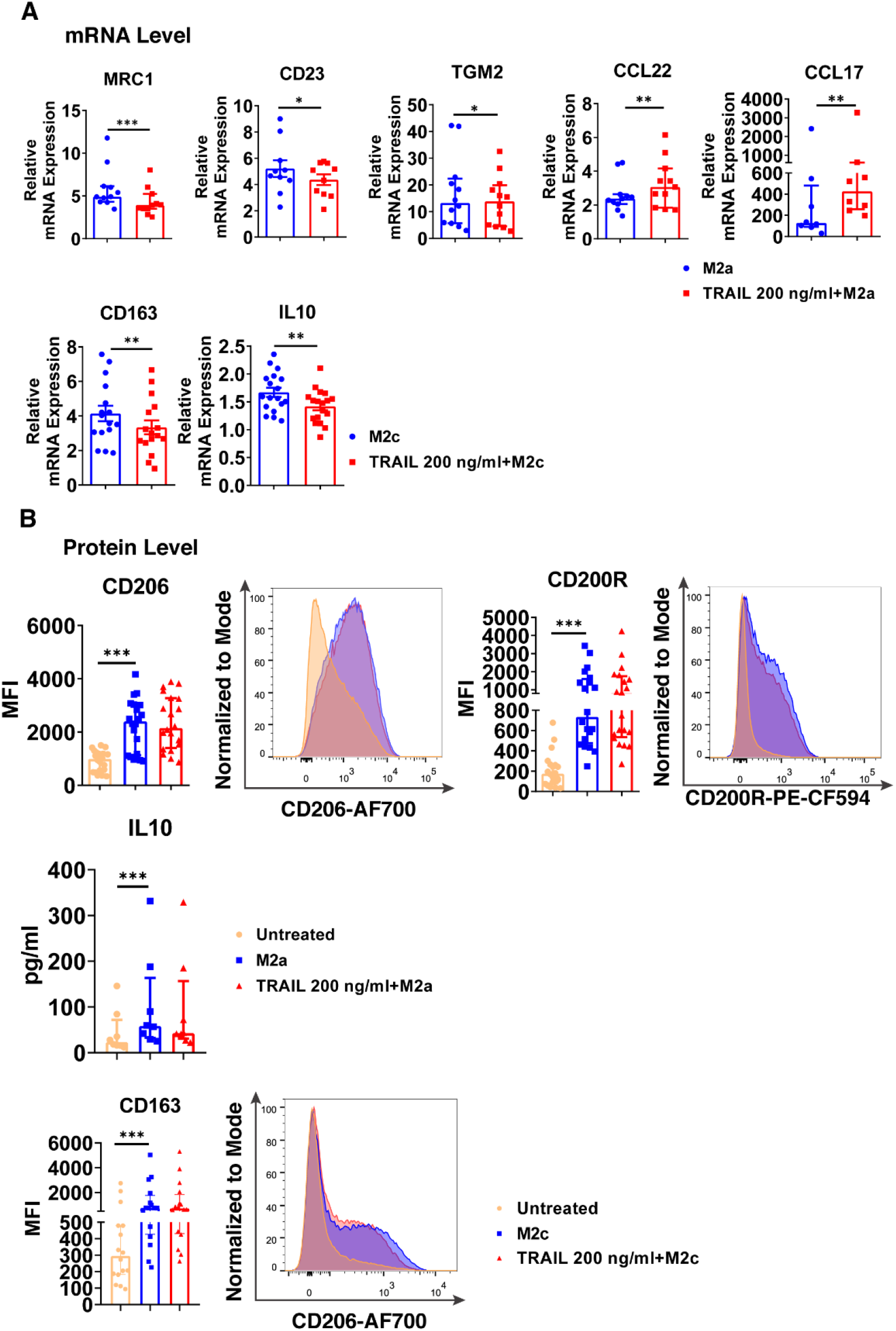
TRAIL changes the expression of the classical M2 markers in primary human M2a and M2c macrophages at the mRNA level. (A) Macrophages were pre-stimulated with 200 ng/ml TRAIL for 2 hours and then polarized into M2a with 20 ng/ml IL-4 or M2c with 10 ng/ml IL-10 for 6 hours. Control group was stimulated only with IL-4 or IL-10 for 6 hours. Expression of M2 markers was analyzed by qPCR. (B) Macrophages were pre-stimulated with 200 ng/ml TRAIL for 6 hours and then polarized into M2a with 20 ng/ml IL-4 or M2c with 10 ng/ml IL-10 for 12 hours. Control groups were left unstimulated or stimulated only with IL-4 or IL-10 for 12 hours. Expression of CD206 and CD200Rin M2a macrophages and CD163 expression in M2c macrophages were analyzed by flow cytometry and representative plots are included. IL-10 production of M2a macrophages was analyzed by ELISA. Data shown are mean ± SEM or median with interquartile range pooled from three or more independent experiments [(A) n=8-18, (B) n= 8-20]. Statistical analyses were performed with a two-tailed paired Student’s t-test, Wilcoxon matched-pairs signed-rank test, or Friedman with Dunn’s multiple comparisons post-hoc test between untreated and M2 macrophages, M2 and TRAIL-treated M2 macrophages, *P<0.05, **P<0.01, ***P<0.001.

In M2c macrophages, TRAIL decreased the expression of the M2 markers CD163 and IL-10 at the mRNA level **(Figure 8A)**; however, no change in the production of the cell surface CD163 was observed at the protein level **(Figure 8B)**.

Furthermore, the production of classical M1 markers was analyzed in M2 macrophages at the protein level. Indeed, TRAIL increased the production of CD86, HLA-DR alpha, and CD64 cell surface activation markers in M2a macrophages, while it increased CD86 and CD64 production in M2c macrophages **(Figure S5)**.

Moreover, the impact of TRAIL on M0 macrophages was investigated. TRAIL increased the expression of classical M1 markers at both mRNA and protein levels **(Figure S1)**; however, it did not significantly affect the expression of M2 markers, except MRC-1 **(Figure S2).**

Our data show that TRAIL affects macrophage polarization by regulating the expression of M1 markers in primary human macrophage subtypes. Overall, our results demonstrate that TRAIL drives macrophages to an M1 phenotype.

### The impact of TRAIL on M2 to M1 switch in primary human macrophages

Tumor cells can direct the associated macrophages into an M2 phenotype, which supports tumor growth and invasion (126, 127, 128). As TRAIL is expressed in the tumor microenvironment (129, 130), the impact of TRAIL on already polarized M2 macrophages was investigated. Primary human macrophages were first polarized into M2a and M2c phenotypes and then treated with TRAIL.

TRAIL increased the expression of intracellular M1 markers at the mRNA level and cell surface activation markers at the protein level in M2a macrophages **(Figure 9)**. Furthermore, a similar trend was observed with TRAIL treatment in M2c macrophages **(Figure 9)**.

**Figure 9.**
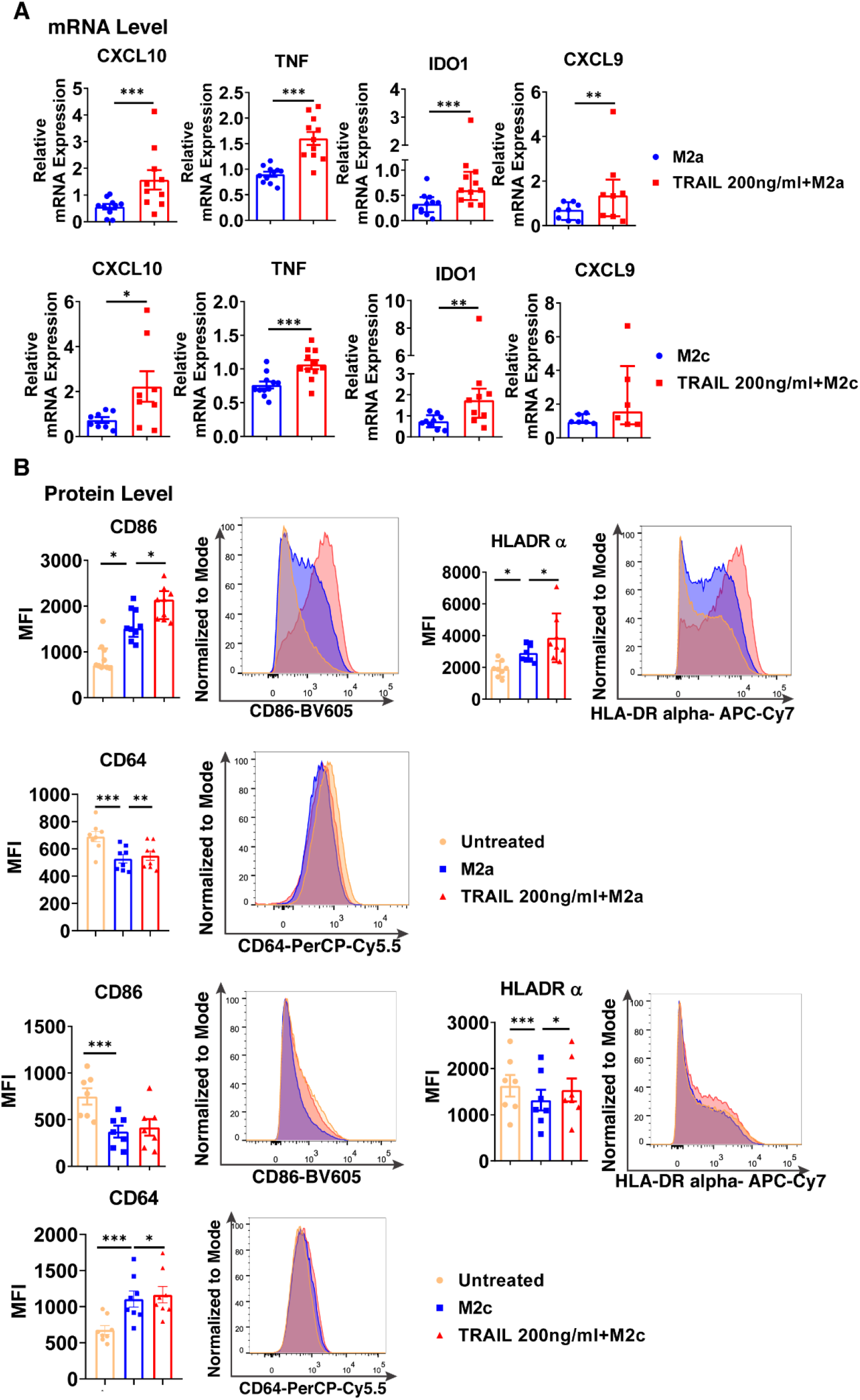
TRAIL shifts primary human M2 macrophages into M1 by upregulating the expression of M1 markers at mRNA and protein levels. (A) Macrophages were polarized into M2a with 20 ng/ml IL-4 or M2c with 10 ng/ml IL-10 for 2 hours and then stimulated with 200 ng/ml TRAIL for 6 hours. Control group was stimulated only with IL-4 or IL-10 for 8 hours. Expression of M1 markers was analyzed by qPCR. (B) Macrophages were polarized into M2a with 20 ng/ml IL-4 or M2c with 10 ng/ml IL-10 for 6 hours and then stimulated with 200 ng/ml TRAIL for 18 hours. Control groups were left unstimulated or stimulated only with IL-4 or IL-10 for 24 hours. Expression of M1 markers was analyzed by flow cytometry and representative plots are included. Data shown are mean ± SEM or median with interquartile range pooled from three or more independent experiments [(A) n=6-11, (B) n=7-9]. Statistical analyses were performed with a two-tailed paired Student’s t-test, Wilcoxon matched-pairs signed-rank test, One-way ANOVA with Sidak’s multiple comparisons post-hoc test, or Friedman with Dunn’s multiple comparisons post-hoc test between untreated and M2 macrophages, M2 and TRAIL-treated M2 macrophages, *P<0.05, **P<0.01, ***P<0.001.

Additionally, the impact of TRAIL on M2-like tumor-associated macrophages (TAMs) was analyzed. In accordance with the literature (131, 132), TAMs were generated *in vitro* by culturing macrophages with different conditioned media (CM) obtained from the AML (U937), bladder (5637) and breast (SKBR3) tumor cell lines and TRAIL was introduced to TAMs without removing the tumor-conditioned media. Subsequently, the changes in the expression of cell surface markers in TRAIL-treated and control group TAMs were analyzed **(Figure S6).**

Consistent with the impact of TRAIL on formerly M2-polarized macrophages **(Figure 9)**, TRAIL treatment increased the expression of cell surface M1 markers in TAMs generated with the CM of three different tumor cell lines **(Figure S6)**. However, TRAIL treatment did not affect the expression of the CD206 M2 marker in TAMs **(Figure S6).**

Therefore, these data show that TRAIL can switch M2 macrophages into a proinflammatory M1 phenotype by upregulating the expression of M1 markers at both mRNA and protein levels.

### The impact of DR4 and DR5 activation on the differentially expressed M1 and M2 markers in primary human macrophages

Next, we investigated through which death receptor TRAIL affects macrophage polarization. Before the analysis of DR4/DR5 activation on macrophages, the distribution of DRs upon polarization and TRAIL treatment was investigated. Human monocyte-derived macrophages express both functional (DR4, DR5) and decoy (DcR1, DcR2) death receptors (34, 59). DRs expression in macrophage subtypes is regulated according to the surrounding milieu. For instance, in the presence of suppressive M2 factors, DR4/DR5 expression is upregulated in human monocyte-derived macrophages (34), THP-1 macrophages, and TAMs compared to M1 (34, 48, 62). On the other hand, in an inflammatory environment such as rheumatoid arthritis or atherosclerosis, DR5 expression is higher in M1 macrophages than in M2 (60, 61, 83, 133).

Intriguingly, contrary to the literature, our result demonstrated that polarization of human macrophages into M1 and M2a decreased cell surface DR5 expression **(Figure S7B-C).** However, no change was observed in the expression of cell surface DR5 with M2c polarization **(Figure S7D).** Regarding DR4 expression, while M1 polarization slightly increased the expression of cell surface DR4, M2c polarization had the opposite impact **(Figure S7B and Figure S7D).** Disparity from the literature may be due to the differences in experimental conditions such as stimulation time (34), or cell type (34, 48, 61, 62, 83, 133).

After the cells were treated with TRAIL, the expression of DR4 was upregulated while DR5 was downregulated at the cell surface in human macrophage subtypes **(Figure S7)**, except for DR4 and DR5 in M1 macrophages **(Figure S7B)** and DR4 in M2c macrophages **(Figure S7D)**.

To investigate which death receptor activation affects differentially expressed M1 and M2 markers, macrophage subtypes were stimulated with DR4 (4C9) or DR5 (D269H/E195R) specific ligands which are generated by changing the receptor specificity of TRAIL by protein engineering method (69, 70, 134). At this stage, cells were stimulated only with DR4 or DR5 ligands or co-treated with DR4 and DR5 ligands simultaneously. Then, the cells were either kept as unpolarized (M0) or polarized into M2 macrophages while control groups were left untreated or treated with only M2 polarization factors. Subsequently, the expression of some of the verified differentially expressed M1/M2 markers was analyzed at the mRNA level in M0, M2a, and M2c macrophages.

DR4 activation did not affect the expression of M1 and M2 markers in M0 macrophages at the mRNA level. On the other hand, DR5 activation mostly increased the expression of M1 markers, while only decreasing the expression of HGF among the M2 markers **(Figure 10A).** Regarding the co-activation of DR4 and DR5 receptors, the expression of all M1 markers was upregulated, while the expression of TMEM37 and HGF was downregulated **(Figure 10A).**

**Figure 10.**
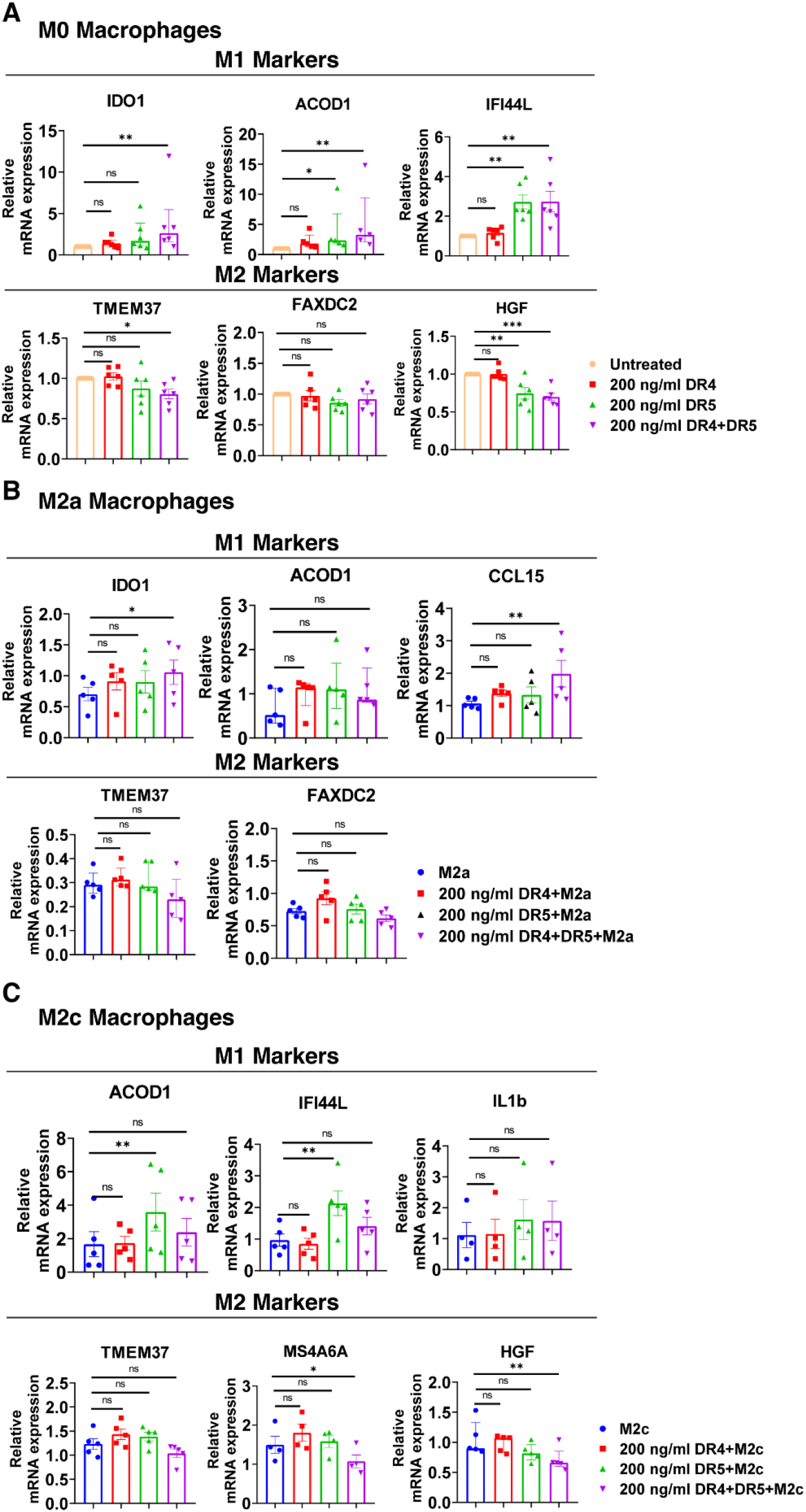
Both DR4 and DR5 receptors regulate primary human macrophage polarization. M0 macrophages were stimulated with 200 ng/ml DR4 or DR5 specific ligands or co-stimulated with DR4 and DR5 specific ligands for 8 hours. For polarization, macrophages were pre-stimulated with only DR4 or only DR5 specific ligands or together with DR4 and DR5 specific ligands for 2 hours and then polarized into M2a (20 ng/ml IL-4) or M2c (10 ng/ml IL-10) for 6 hours. Control groups were left unstimulated or stimulated with only M2a or M2c polarization factors for 6 hours. qPCR analyses of M1 and M2 markers in (A) M0, (B) M2a, and (C) M2c macrophage subtypes were shown. Data shown are mean ± SEM or median with interquartile range pooled from two independent experiments [(A) n=5-6, (B) n=5, (C) n=4-5]. Statistical analyses were performed with a One-way ANOVA with Dunnet’s multiple comparisons post-hoc test, or Friedman with Dunn’s multiple comparisons post-hoc test between control and treated groups, *P<0.05, **P<0.01, ***P<0.001.

In M2a macrophages, only the co-activation of DR4 and DR5 increased the expression of M1 markers IDO1 and CCL15 at the mRNA level, while it did not affect the expression of M2 markers (**Figure 10B).** In M2c macrophages, as observed in M0, DR5 activation was required to increase the expression of M1 markers at the mRNA level. However, to decrease the expression of M2 markers MS4A6A and HGF, co-stimulation of both DR4 and DR5 was needed **(Figure 10C).** These results show that both DR4 and DR5 receptors play a role in primary human macrophage polarization; however, the regulation of M1/M2 markers expression mostly depends on DR5-mediated activation.

### The impact of TRAIL on macrophage cytotoxicity against acute myeloid leukemia (AML)

Acute myeloid leukemia (AML), occurring due to the abnormal proliferation of myeloid progenitor cells in the bone marrow, comprises the majority of acute leukemia diagnoses (135, 136, 137). Studies showed that the frequency of tumor-promoting M2 macrophages (CD163+ CD206+) in the bone marrow of AML patients is elevated compared to healthy donors (138) which decreases patient survival (139). Therefore, targeting macrophage polarization could be an effective way of the treatment of AML patients. To investigate the impact of TRAIL, a widely used agent in cancer clinical studies (140, 141) on macrophage cytotoxic response against AML cells, macrophages were co-cultured with the acute myeloid leukemia (AML) cell line (U937) and macrophage-induced AML cell killing was determined.

M1 macrophages can contribute to the anti-tumorigenic response via a variety of mechanisms, involving the production of pro-inflammatory cytokines such as TNF (122, 142, 143, 144), reactive oxygen species (143, 144, 145), and antimicrobial peptides (145). Besides, M1 macrophages can eliminate tumor cells by phagocytosis (146, 147).

TRAIL treatment significantly enhanced the cytotoxicity of M1 macrophages against the U937 cells and increased tumor cell death compared to M1 macrophages alone **(Figure 11).** Figure 11 shows that there is a donor-to-donor variability in macrophage cytotoxicity. Some donors had high M1 cytotoxicity against cancer cells in their control groups and TRAIL increased macrophage cytotoxicity in these donors. However, in the donors who had low M1 cytotoxicity in the control groups, TRAIL did not make a major impact.

**Figure 11.**
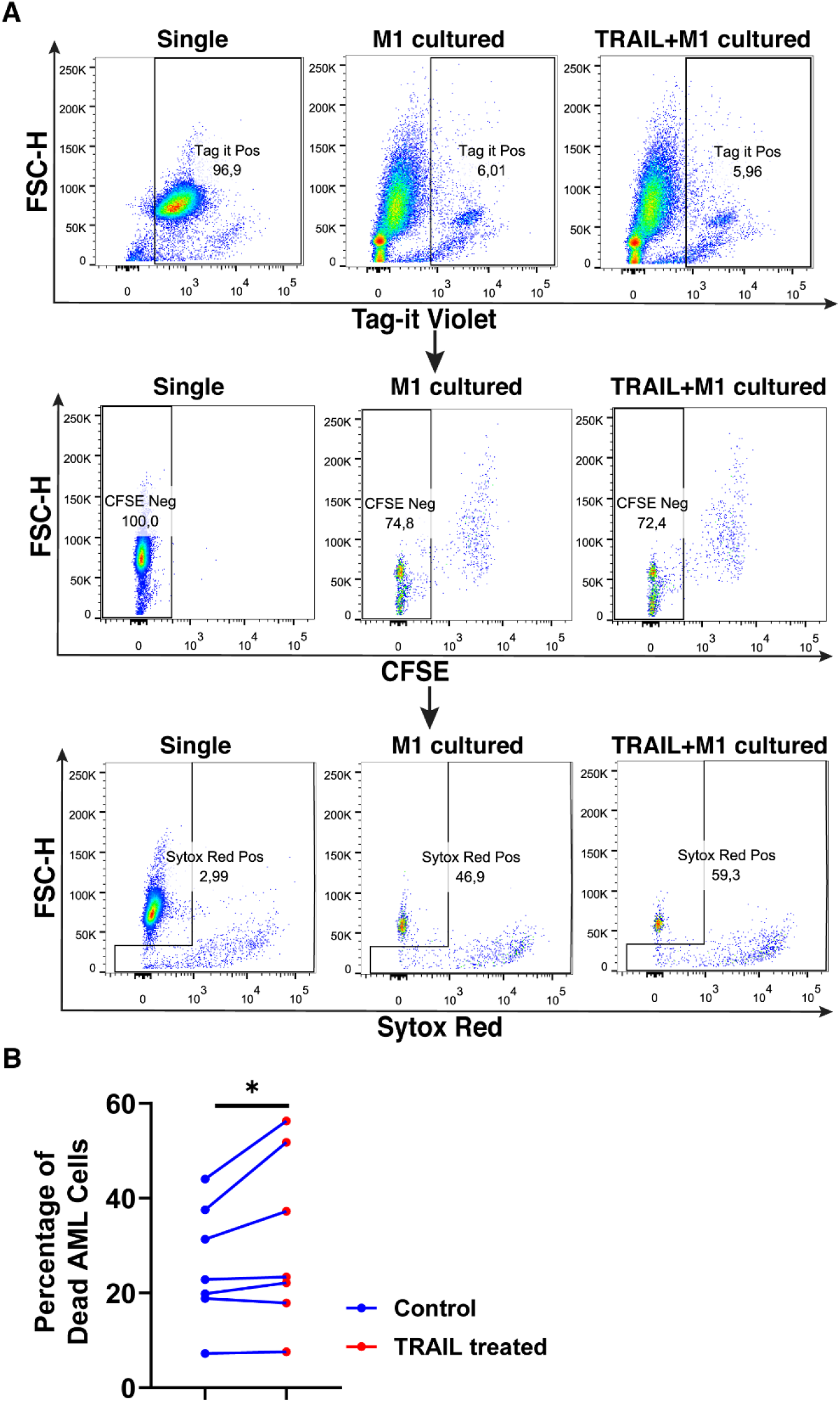
TRAIL increases macrophage cytotoxicity against AML cells. Macrophages were labelled with the cell tracker CFSE. Control and TRAIL-pre-stimulated (200 ng/ml, 6 h) macrophages were polarized into M1 with 100 ng/ml LPS+20 ng/ml IFNγ for 12 hours. After polarization, macrophages were washed and co-cultured with Tag-IT cell tracker labelled U937 cells for 72 hours. Tag it-Violet^+^ CFSE^-^ tumor cell death was assessed with Sytox Red staining and analyzed with flow cytometry. Macrophage-mediated tumor cell killing (specific cytotoxicity) was calculated as the percentage of Sytox Red^+^ AML cells (Tag-IT Violet^+^ CFSE^-^) co-cultured with macrophages (CFSE^+^)-the percentage of Sytox Red^+^ AML cells alone. (A) Representative dot plots showing the gating strategy and (B) bar graphs showing specific cytotoxicity (Tag-IT Violet^+^ CFSE^-^ Sytox Red^+^). Data shown as symbols and lines pooled from four independent experiments (n= 7). Statistical analyses were performed with a two-tailed paired Student’s t-test, between the control and TRAIL-treated groups, *P<0.05.

The impact of TRAIL on the phagocytotic activity of macrophages was also analyzed **(Figure 12).** In accordance with the impact on cytotoxicity against tumor cells, TRAIL stimulation slightly increased the phagocytosis of tumor cells by M1 macrophages **(Figure 12).** These results show that even though the donor variability is high, overall, TRAIL treatment moderately enhances both phagocytosis of AML cells and cytotoxic cancer cell killing by macrophages.

**Figure 12.**
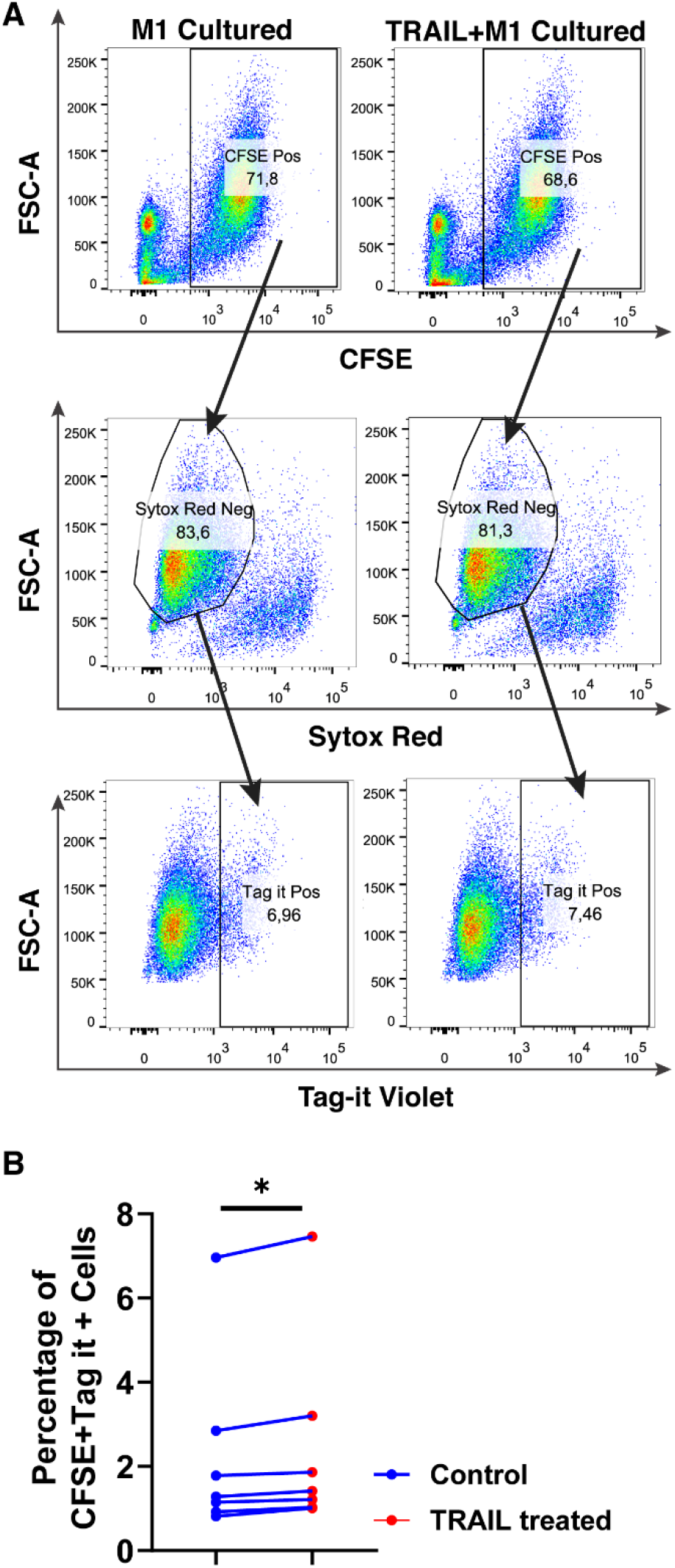
TRAIL slightly increases phagocytosis of AML cells by macrophages. Macrophages were labelled with the cell tracker CFSE. Control and TRAIL-pre-stimulated (200 ng/ml, 6 h) macrophages were polarized into M1 with 100 ng/ml LPS+20 ng/ml IFNγ for 12 hours. After polarization, macrophages were washed and co-cultured with Tag-IT labelled U937 cells for 72 hours. Tag-IT Violet^+^ AML cells within live macrophages (CFSE^+^ Sytox Red^-^) were analyzed by flow cytometry. (A) Representative dot plots with a gating strategy and (B) Bar graphs show the percentage of double positive live macrophages (CFSE^+^ Sytox Red^-^ Tag-IT Violet^+^). Data shown as symbols and lines pooled from four independent experiments (n=7). Statistical analyses were performed with Wilcoxon matched-pairs signed-rank test between control and TRAIL-treated groups, *P<0.05.

### The impact of TRAIL on the clinical outcome of cancer patients

TRAIL expression in the tumor microenvironment (TME) is highly encountered in various tumor types (129, 130). After observing that TRAIL increases M1 macrophage cytotoxicity against tumor cells, the effect of TRAIL on the clinical outcome of cancer patients was investigated by analyzing KM plotter and cBioPortal databases. First, cancer patients were stratified based on the median expression of TRAIL, and then the association between the level of TRAIL expression and the overall survival of patients was analyzed. Patients with high TRAIL expression had longer overall survival in sarcoma cancer but this correlation was not observed in ovarian cancer patients **(Figure 13A)**. However, in both cancer types, when patients were dichotomized into macrophage-rich versus macrophage-low groups, high TRAIL expression positively correlated with longer overall survival of patients in the cases with high tumor macrophage content, but not in low macrophage content. **(Figure 13A)**. Furthermore, TRAIL expression had a positive correlation with M1-related gene signature (CXCL10, CXCL11, IFI44L, CD38) while having no or negative correlation with M2-related gene signature (TGM2, HGF, HPGD, FAXDC2) in ovarian and sarcoma cancer patients **(Figure 13B)**. These results show that TRAIL expression is positively correlated with increased expression of M1 markers in tumors from ovarian and sarcoma patients and longer survival in cases with high, but not low, tumor macrophage content.

**Figure 13.**
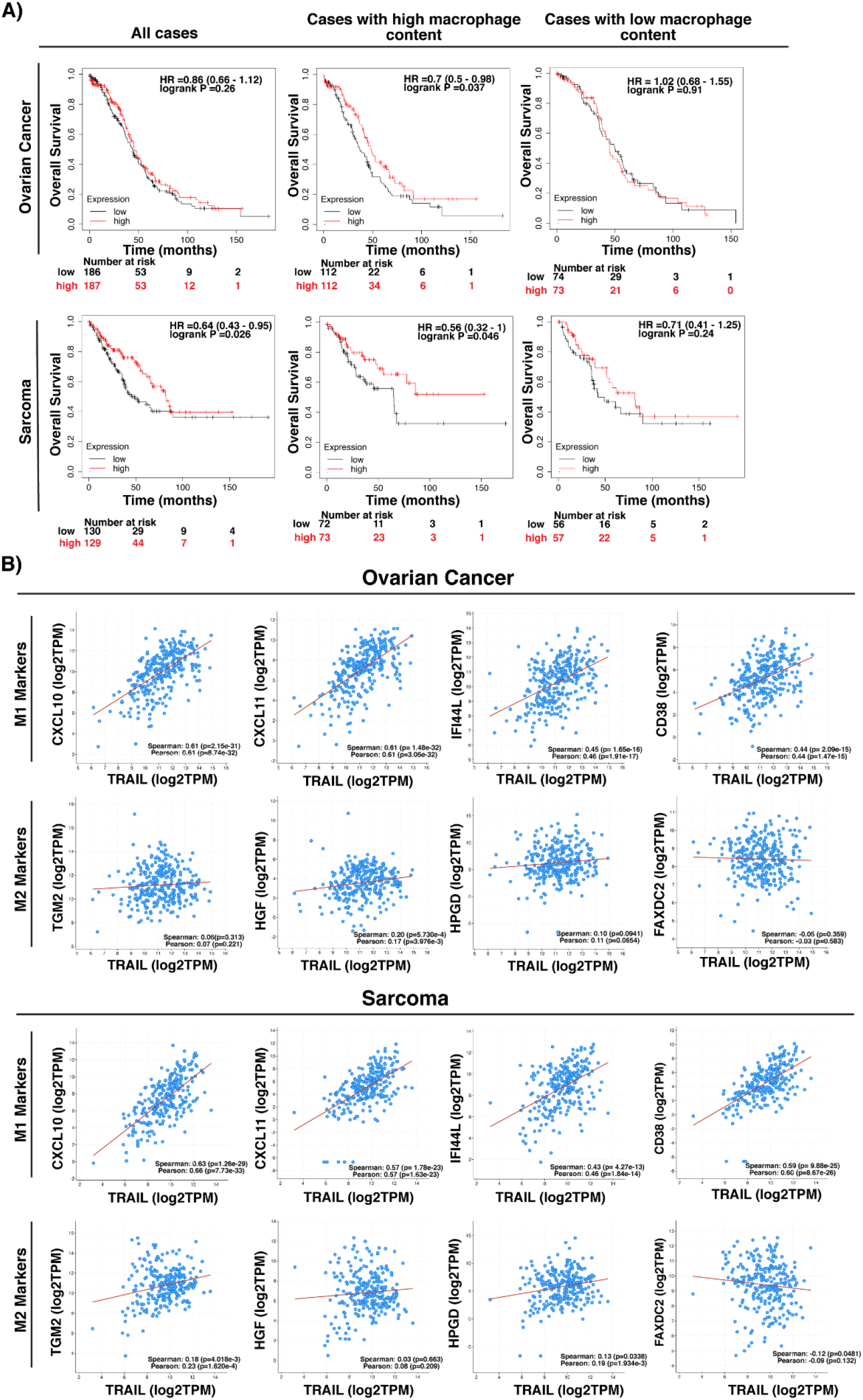
High-level expression of TRAIL in ovarian and sarcoma cancer patients is positively correlated with M1 gene signature and increased survival in cases with high tumor macrophage content. (A) Kaplan-Meier plots showing the overall survival of ovarian and sarcoma cancer patients based on TRAIL gene (TNFSF10) expression. Patients were classified as either high (red line) or low (black line) TRAIL expression based on the median cut-off value. Hazard ratio (HR), p-value, and confidence interval (CI) for each group were determined using log-rank test. (B) Scatter plots showing the correlation between the expression of TRAIL and M1 markers (CXCL10, CXCL11, IFI44L, and CD38) or M2 markers (TGM2, HGF, HPGD, and FAXDC2) in ovarian and sarcoma cancer patients. Spearman’s rank and Pearson correlation coefficient and p-values were also shown. A positive correlation (R>0) indicates that higher expression of TRAIL is associated with higher expression of the M1 markers at the gene level. The plots were generated by using KM plotter and cBioPortal tools.

In addition to transcriptomic analysis, proteomic analysis was also performed in cancer patients by using Clinical Proteomic Tumor Analysis Consortium (CPTAC) database (https://cptac-data-portal.georgetown.edu/cptacPublic/) to demonstrate the association of TRAIL presence with the expression of M1 and M2 markers at protein level. Our data showed that in ovarian cancer patients, TRAIL expression was positively correlated with the expression of M1 markers (CD38 and IFI44L) while having no correlation with M2 markers (TGM2 and HPGD) at the protein level **(Figure 14).** Overall, the results obtained from the analysis of transcriptomic (TCGA) and proteomic (CPTAC) datasets show that the presence of TRAIL in the tumor microenvironment is positively correlated with M1 signature **(Figure 13 and Figure 14).**

**Figure 14.**
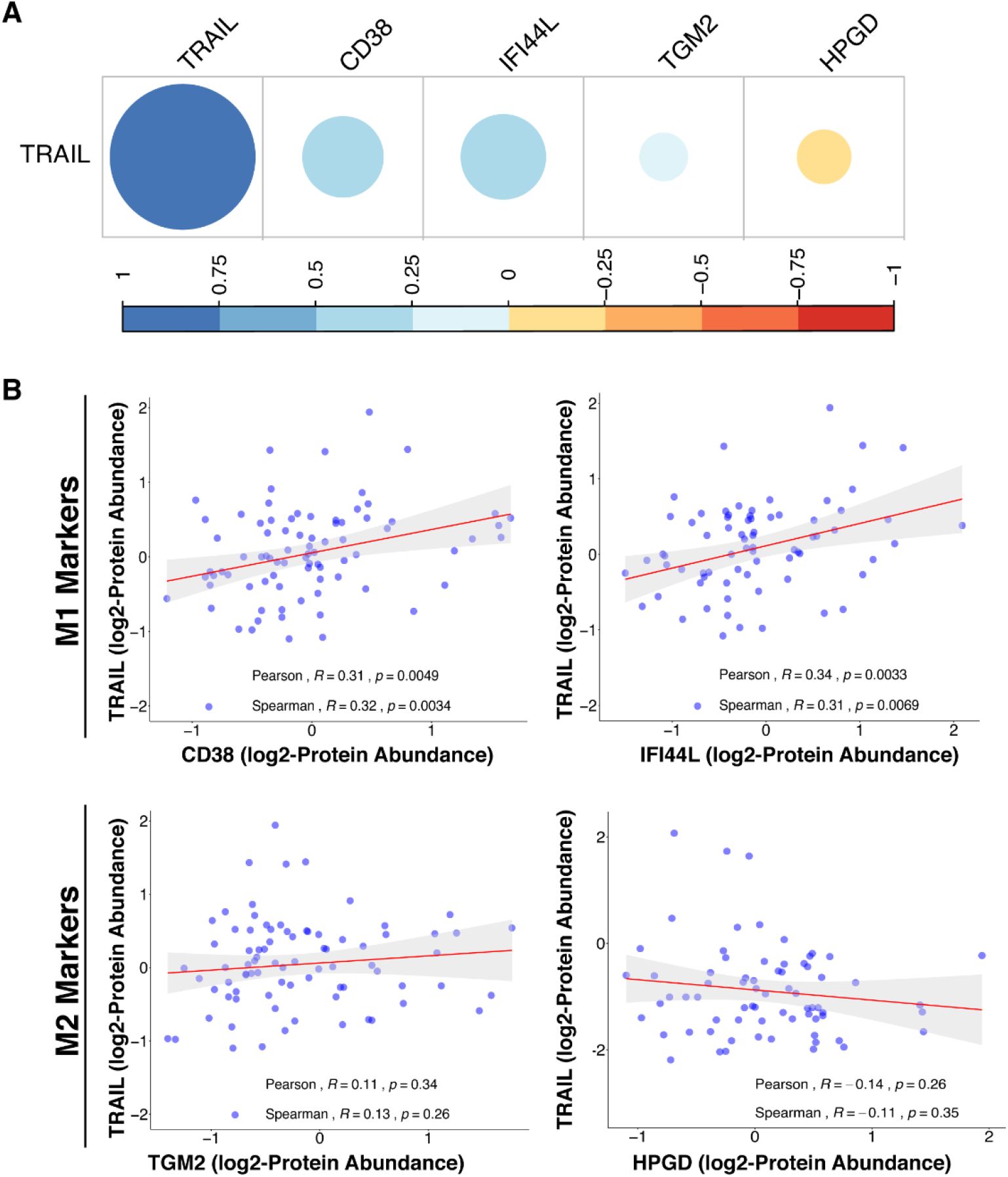
TRAIL expression is positively correlated with M1 proteomic signature in ovarian cancer patients. (A) The correlation matrix between the expression of TRAIL and M1 (CD38 and IFI44L) or M2 (TGM2 and HPGD) markers in ovarian cancer patients was shown based on the correlation coeffient values. (B) Scatter plots showing the correlation between the expression of TRAIL and each M1 or M2 marker were demonstrated with Spearman’s rank and Pearson correlation coefficient and p-values. A positive correlation (R>0) indicates that higher expression of TRAIL is related with higher expression of the M1 markers at the protein level.

## DISCUSSION

Macrophages can be polarized into two main subtypes: pro-inflammatory/anti-tumorigenic M1 macrophages and anti-inflammatory/tumor-promoting M2 macrophages (3, 4) in the presence of certain polarization factors. Disruption of the balance between M1 and M2 macrophages can cause progression of various types of diseases. For instance, obesity and type II diabetes are associated with an increased number of M1 macrophages; while allergic asthma and cancer are related to an increased number of M2 macrophages (12, 17, 148). Therefore, identifying modulators regulating macrophage polarization is important for the design of effective macrophage-mediated immunotherapies.

TRAIL mainly provides homeostasis by eliminating activated effector immune cells and inducing selective toxicity in transformed cells (47, 48, 65, 149, 150, 151, 152); however, it has been reported that the presence of TRAIL can exacerbate inflammation as well (153, 154, 155, 156). Since tumor cells are more susceptible to TRAIL-induced cell death compared to healthy cells, and TRAIL can selectively target tumor-associated feeder cells, TRAIL has been the center of cancer therapy studies (22, 24, 45, 46, 47, 48, 157).

Studies to date in macrophages generally focus on the effect of TRAIL on macrophage survival and have shown that the sensitivity of macrophage phenotypes to TRAIL can vary depending on the microenvironment (34, 47, 48, 60, 61, 62). Apart from this, one study has reported that TRAIL also triggers survival pathways in primary human and mouse macrophages and induces the production of TNF, IL-1β, and IL-6 pro-inflammatory cytokines (63). However, the effect of TRAIL on primary human macrophage polarization and whether it can be used as a modulator in macrophage response has not been investigated.

The soluble TRAIL used in the study did not majorly affect the viability of primary human macrophage subtypes **(Figure 1)** which conflicts with the literature. This suggests that usage of different types of TRAIL formulations such as Recombinant Super Killer TRAIL, which has an enhanced level of cross-linkage among others (34), or agonistic antibodies targeting DR5 (TRA-8) (60) can initiate a stronger impact on cell death compared to soluble TRAIL ligand.

RNA sequencing and qPCR analyses depicted that TRAIL increased the expression of M1 markers while decreasing the expression of M2 markers at the mRNA level in both M0 and M2 (M2a, M2c) polarized macrophage subtypes **(Figure 2**, **Figure 3, Table S1, Figure S1A)**. Regarding M1 macrophages, although upon TRAIL treatment there was a slight increase in the expression of the M1 markers CXCL10, CXCL11, and IDO1, and a slight decrease in the expression of the M2 markers TMEM37 and ANGPTL4, this did not reach statistical significance **(Table S1B)**. This could be due to the variation in gene expression in primary human samples from different donors (158). On the other hand, the qPCR analysis showed that TRAIL increased the expression of the classical M1 markers in M1 macrophages **(Figure 7A)**. Furthermore, TRAIL reduced the expression of classical M2 markers in M2 macrophages at the mRNA level **(Figure 8A)**. In brief, TRAIL successfully enhances M1 response by regulating the expression of both classical and newly determined M1/M2 markers at the mRNA level in human macrophage subtypes.

Concerning the impact of TRAIL at the protein level, it mostly increased the expression of both classical and newly determined M1 markers in all macrophage subtypes (M0, M1, M2a, and M2c) **(Figure 4-6**, **Figure 7B, Figure S1B, Figure S3, Figure S5)**; however, it did not change the production of the M2 markers in M2 macrophages **(Figure 8B, Figure S2B).** While TRAIL can upregulate the expression of M1 markers in human macrophage subtypes at both gene and protein levels, this effect was not observed in M2 markers at the protein level even though established downregulation at the mRNA level. The expressions at mRNA and protein levels do not always show a direct one-to-one correlation (35). The translational rate (159, 160, 161) and post-translational modifications (102, 162) also regulate the production/stability of the proteins which might interfere with the effect of TRAIL on the expression of the M2 markers. In accordance with our results, previous studies reported that a modulator can direct macrophages into an M1 phenotype without any change in the expression of M2 markers at the protein level (163, 164).

The most prominent effect of TRAIL on M1 macrophages was increasing the production of M1 chemokine CXCL10 and the cytokines IL-1β, TNF, and IL-12p70 **(Figure 7B).** Furthermore, increased expression of surface activation markers CD86, HLA-DR alpha, and CD64 **(Figure 7B)** in M1 macrophages suggests that TRAIL could support the anti-tumorigenic response via providing cell-to-cell contact and recruiting/generating effector immune cells such as NK, DC, Th1, and Th17 cells (5, 165, 166, 167, 168). Regarding M2a macrophages, TRAIL treatment upregulated the expression of the M2 chemokines CCL22 and CCL17. Although these chemokines are induced by M2a stimulants, the upregulation of their expression by inflammatory agents such as LPS (169), their association with the progression of autoimmune diseases such as arthritis (170, 171), and their role in attracting the anti-tumorigenic Th17 cells (166, 167, 172, 173) suggest that TRAIL may enhance inflammatory response in M2a macrophages by increasing the expression of these chemokines. Furthermore, TRAIL increased the expression of M1 surface activation markers in M2 macrophage subtypes at the protein level **(Figure S5)**, which are associated with antigen presentation/co-stimulation (CD86 and HLA-DR) (174) and production of ROS and inflammatory cytokines (CD64) (175, 176, 177, 178, 179). Although these surface molecules are expressed in M2 macrophages, they are not expressed as highly as in M1 macrophages (177, 180, 181). M1 macrophages are more potent in antigen presentation (174) and the production of ROS (177) by using these receptors. Therefore, TRAIL may also contribute to the anti-tumorigenic response by enhancing antigen presentation/co-stimulation, and cytotoxic function of M2 macrophages.

As demonstrated in Figure 10, both DR4 and DR5 receptors regulate human macrophage polarization; however, the regulation of M1/M2 markers expression mostly depends on DR5-mediated activation. Besides, apart from DR4 and DR5, the DcR2 decoy receptor, that is shown to induce survival pathways in cancer cells (27, 182) and expressed by human macrophages (34, 59), might also play a role in TRAIL-mediated macrophage polarization. To investigate this, DR4/DR5 co-stimulation together with DcR2 activation or TRAIL stimulation in the presence of DcR2 neutralization could be tested to determine its role in primary human macrophage polarization. Apart from these, the interaction of the TRAIL ligand to target death receptors may differ from the individual DR4/5 stimulation. In that case, a structural analysis could be performed to have a deeper understanding about the binding of TRAIL vs DR4/DR5 ligands to target receptor/s. In summary, although mainly DR5 appears to play a role in macrophage polarization, the regulation of each M1/M2 marker differs from one another and could require activation of different receptors.

Our study showed that TRAIL also increased the expression of M1 markers in formerly M2-polarized macrophages and TAMs generated by tumor-conditioned medium **(Figure 9, Figure S6).** This impact is quite critical in the tumor environment since directing macrophages into an M1 phenotype prevents tumor progression (5, 123, 126, 127, 128). Addional experiments could be performed with TAMs sorted from patients tumor samples in follow-up studies. In the co-culture system, macrophage cytotoxicity against the U937 AML cell line was slightly enhanced with TRAIL stimulation **(Figure 11**, **Figure 12).** Although TRAIL did not very potently increase macrophage cytotoxicity in all donors, there was a prominent increase in the donors which showed higher cytotoxic response against tumor cells. This shows that donor responses could be quite different in primary human samples and TRAIL only enhances the tumor cell killing ability of the macrophages which are good responders.

The low-level phagocytosis of AML cells by macrophages including both control and TRAIL-treated groups suggests that either this anti-tumorigenic response is not used by macrophages against U937 AML cell line or TRAIL treatment is not sufficient. Therefore, CD47, which is highly expressed in AML cells and hinders phagocytosis via interacting with SIRP1α on target cells (183, 184), can be neutralized by using a monoclonal antibody and the synergistic impact of TRAIL and anti-CD47 antibody on macrophage phagocytosis of tumor cells can be investigated for further studies. Overall, TRAIL stimulation increased tumor cytotoxicity of macrophages to some extent. To provide a more comprehensive conclusion to the anti-tumorigenic role of TRAIL, alternative soft/solid tumors could be utilized, or the indirect effect of TRAIL on the other immune effector cells (NK cells, iNKT cells, and T cells) through macrophage activation could also be analyzed. Involving other effector cells in the co-culture may induce a more potent tumor cytotoxicity response than macrophages alone. In addition, Do-Thi et al. reported that an exogenous modulator can enhance the anti-tumorigenic response of macrophages without affecting phagocytosis or cytotoxicity functions but inducing the production of chemokines that recruits effector cells into the tumor microenvironment (164). Similar to this study, since TRAIL increases the production of anti-tumorigenic chemokines in macrophages, the impact of macrophage-mediated chemokines on the migration of other effector cells can also be performed in future studies. Clinical data analyses were also carried out to assess the clinical relevance of our findings in cancer patients **(Figure 13 and Figure 14).** In this regard, it was observed that in ovarian and sarcoma cancer patients, TRAIL expression was positively correlated with the expression of M1 markers in the tumor microenvironment. In addition, high TRAIL expression was positively correlated with increased survival in ovarian and sarcoma cancer patients with high, but not low, tumor macrophage content. TRAIL expression was also positively correlated with M1 related proteomic signature in ovarian cancer patients. This suggests that the presence of TRAIL could impact the survival of patients by converting macrophages into M1 in certain cancer types. To clarify these points, the correlation between TRAIL expression and the levels of M1 and M2 macrophages in the tumors from cancer patients could be analyzed *ex vivo* in follow-up studies.

In conclusion, our findings show that TRAIL polarizes primary human macrophages into pro-inflammatory M1 phenotype by affecting the production of M1 markers rather than M2 markers. Our study sheds new light onto mechanisms of macrophage polarization by defining TRAIL as a new regulator. TRAIL is investigated as a therapeutic agent in clinical trials due to its selective toxicity to transformed cells. Our study suggests that TRAIL could also enhance anti-tumorigenic response by directing primary human macrophage subtypes into an M1 phenotype.

## Data availability statement

The data presented in the study are deposited in the NCBI GEO database with the accession number GSE227737.

## Ethics statement

The study was reviewed and approved by the Non-Invasive Research Ethical Committee of Dokuz Eylul University (Approval number: 2018/06-26) and the University of Galway, Ireland Research Ethics Committee (Approval number: 2022.02.022) for the use of buffy coats. The buffy coats were obtained from healthy donors who provided their written informed consent at Dokuz Eylul University Blood Bank (Izmir, Turkey) and University of Galway, Ireland.

## Authors contributions

SG and DGH performed the experiments and analyzed the data. GK, AO, AB, AC, HG and ES performed the bioinformatic analysis. DS, SG, DGH and ES contributed to the design and conception of the study. DS, SG, DGH, GK, AO and ES wrote the manuscript. DS supervised the whole study. All authors contributed to the study and approved the submitted version.

## Acknowledgments

We thank Dr. Yavuz Dogan and the laboratory staff at Dokuz Eylul University Blood Bank for supplying buffy coats and for their kind assistance. We thank Dr. Esra Erdal and Dr. Serap Erkek for providing the tumor cell lines. We also thank the staff of the Flow Cytometry and Cell Sorting Facility at the Izmir Biomedicine and Genome Center and Apoptosis Research Center, University of Galway, Ireland for their technical support. We thank Dr. Asli Suner Karakülah for the statistics consultation. We also thank, Duygu Unuvar, Asli Korkmaz and Mark Gurney for their valuable support.

## Publisher’s note

All claims expressed in this article are solely those of the authors and do not necessarily represent those of their affiliated organizations, or those of the publisher, the editors, and the reviewers. Any product that may be evaluated in this article, or claim that may be made by its manufacturer, is not guaranteed or endorsed by the publisher.

## Supplementary material

The Supplementary Material for this article can be found online at:

