## Supplementary Tables and Figures for "TRAIL promotes the polarization of human macrophages toward a proinflammatory M1 phenotype and is associated with increased survival in cancer patients with high tumor macrophage content"

#### Supplementary Data

#### Supplementary Tables

**Table S1. FDR and fold change values of differentially expressed M1 and M2 markers in primary human M0, M1, M2a, and M2c macrophages upon TRAIL treatment.** According to RNA sequencing analysis, TRAIL-induced differentially expressed M1 and M2 markers in (A) M0, (B) M1, (C) M2a, and (D) M2c macrophage subtypes were determined (n=3-4). FDR values and fold changes of markers were depicted as tables. **(Supplement of Figure 2)**

**A**

**M0 Macrophages**

| M1 Genes | LogFC | LogCPM | LR | P-Value | FDR | Fold Change |
|---|---|---|---|---|---|---|
| CCL15 | 2,907937 | -1,274537 | 38,70564 | 4,93E-10 | 9,62E-08 | 7,505439771 |
| CXCL11 | 2,734056 | 0,0373049 | 68,3073 | 1,4E-16 | 9,47E-14 | 6,65323491 |
| CXCL10 | 2,608033 | 2,530077 | 29,67207 | 5,12E-08 | 5,89E-06 | 6,096720046 |
| IFI44L | 2,087466 | 3,8332614 | 64,71575 | 8,65E-16 | 4,82E-13 | 4,250008526 |
| CXCL1 | 1,874095 | 2,9751543 | 71,44037 | 2,86E-17 | 2,39E-14 | 3,665715319 |
| IL12B | 1,814687 | -2,129927 | 12,88084 | 0,000332 | 0,01117 | 3,517831879 |
| IDO1 | 1,682171 | 1,9116785 | 46,62585 | 8,59E-12 | 2,45E-09 | 3,209104641 |
| ACOD1 | 1,669245 | -0,286877 | 29,94927 | 4,43E-08 | 5,3E-06 | 3,18048193 |
| CD38 | 1,374008 | 2,7878062 | 68,28464 | 1,42E-16 | 9,47E-14 | 2,591896568 |
| IL1B | 0,747764 | 3,5410055 | 4,33445 | 0,037348 | 0,248974 | 1,679188778 |
| **M2 Genes** | **LogFC** | **LogCPM** | **LR** | **P-Value** | **FDR** | **Fold Change** |
| SELENOP | -1,59326 | 4,6101703 | 24,93365 | 5,93E-07 | 5,51E-05 | -3,017304424 |
| HGF | -1,49268 | 4,2914729 | 29,08246 | 6,94E-08 | 7,73E-06 | -2,814112767 |
| F13A1 | -1,42713 | -0,291227 | 9,979399 | 0,001583 | 0,035955 | -2,68911441 |
| FAXDC2 | -1,27713 | 4,1928475 | 12,88344 | 0,000332 | 0,01117 | -2,423565133 |
| MS4A6A | -1,16267 | 5,6874632 | 17,60946 | 2,71E-05 | 0,001531 | -2,238707242 |
| TMEM37 | -1,09938 | 4,2562729 | 42,63845 | 6,59E-11 | 1,49E-08 | -2,142629269 |
| HTR2B | -0,47738 | 1,9504305 | 3,053927 | 0,080542 | 0,363538 | -1,3922159 |
| PRR5L | -0,26564 | 2,9720773 | 3,074031 | 0,079552 | 0,360884 | -1,202171658 |
| ANGPTL4 | -0,14318 | -0,144882 | 0,212906 | 0,644499 | 0,869164 | -1,104333412 |
| HPGD | -0,139 | 3,5830761 | 1,329761 | 0,248848 | 0,600809 | -1,101141603 |

**B**

**M1 Macrophages**

| **M1 Genes** | **LogFC** | **LogCPM** | **LR** | **P-Value** | **FDR** | **Fold Change** |
|---|---|---|---|---|---|---|
| CXCL10 | 0,828947 | 11,228708 | 2,327726 | 0,127087 | 0,965829 | 1,77638816 |
| CXCL11 | 0,822665 | 9,8807491 | 1,64701 | 0,199366 | 0,965829 | 1,768670619 |
| IDO1 | 0,68372 | 10,636941 | 2,06094 | 0,151117 | 0,965829 | 1,606276149 |
| CD38 | 0,449123 | 8,7203642 | 3,482029 | 0,062039 | 0,965829 | 1,365210544 |
| IFI44L | 0,385988 | 8,7063798 | 3,474881 | 0,062307 | 0,965829 | 1,306754076 |
| ACOD1 | 0,155129 | 10,641016 | 0,993177 | 0,318967 | 0,965829 | 1,113521407 |
| CCL15 | 0,0036 | 4,2821061 | 0,000529 | 0,981656 | 1 | 1,002498528 |
| CXCL1 | -0,15699 | 5,8439745 | 0,650235 | 0,420029 | 0,965829 | -1,114960656 |
| IL12B | -0,35411 | 9,9254864 | 0,925849 | 0,335944 | 0,965829 | -1,278193567 |
| IL1B | -0,44167 | 11,140957 | 1,828917 | 0,176256 | 0,965829 | -1,358174659 |
| **M2 Genes** | **LogFC** | **LogCPM** | **LR** | **P-Value** | **FDR** | **Fold Change** |
| TMEM37 | -1,32049 | -1,105387 | 3,747579 | 0,052884 | 0,965829 | -2,49750808 |
| ANGPTL4 | -0,98658 | 3,1596824 | 3,76906 | 0,052209 | 0,965829 | -1,981485499 |
| HPGD | -0,48634 | 2,1677845 | 1,002175 | 0,316785 | 0,965829 | -1,400887258 |
| FAXDC2 | -0,29119 | 0,6192352 | 0,637408 | 0,424651 | 0,965829 | -1,22365007 |
| MS4A6A | -0,23175 | 3,8016707 | 1,266893 | 0,26035 | 0,965829 | -1,174258843 |
| HTR2B | -0,08973 | -1,410394 | 0,026217 | 0,871373 | 0,995379 | -1,064168801 |
| SELENOP | -0,05307 | 2,1319047 | 0,053723 | 0,816707 | 0,989967 | -1,037471183 |
| F13A1 | 0,194691 | -1,682 | 0,111502 | 0,73844 | 0,982077 | 1,144478827 |
| PRR5L | 0,137365 | 0,3953011 | 0,14339 | 0,704934 | 0,981564 | 1,099894234 |
| HGF | 0,119124 | 0,7295354 | 0,178002 | 0,673096 | 0,977811 | 1,086075062 |

C

#### M2a Macrophages

| M1 Genes | LogFC | LogCPM | LR | P-Value | FDR | Fold Change |
|---|---|---|---|---|---|---|
| CXCL10 | 2,846259 | 0,2848345 | 7,784432 | 0,00527 | 0,041815 | 7,191330785 |
| IL12B | 2,787451 | -3,782405 | 7,530861 | 0,006065 | 0,046259 | 6,904087519 |
| IFI44L | 1,980685 | 2,1138425 | 203,2681 | 4,04E-46 | 4,51E-43 | 3,946803029 |
| ACOD1 | 1,923619 | -2,275701 | 17,71986 | 2,56E-05 | 0,000531 | 3,793734083 |
| CXCL1 | 1,691321 | -0,956927 | 32,14145 | 1,43E-08 | 6,29E-07 | 3,229522316 |
| IL1B | 1,597582 | 1,9860465 | 8,996606 | 0,002705 | 0,025411 | 3,026355753 |
| CXCL11 | 1,581527 | -1,267195 | 18,66776 | 1,56E-05 | 0,000347 | 2,992865402 |
| IDO1 | 1,502411 | -0,291499 | 38,79287 | 4,71E-10 | 2,79E-08 | 2,833158513 |
| CCL15 | 1,278511 | -0,849975 | 17,46876 | 2,92E-05 | 0,000595 | 2,42588452 |
| CD38 | 1,152447 | 1,5231751 | 71,05027 | 3,48E-17 | 5,42E-15 | 2,222906326 |
| **M2 Genes** | **LogFC** | **LogCPM** | **LR** | **P-Value** | **FDR** | **Fold Change** |
| TMEM37 | -0,94794 | 2,691378 | 68,89079 | 1,04E-16 | 1,55E-14 | -1,929122046 |
| HTR2B | -0,91003 | -0,673713 | 10,71438 | 0,001063 | 0,012124 | -1,87908426 |
| FAXDC2 | -0,60759 | 3,2927398 | 45,33441 | 1,66E-11 | 1,29E-09 | -1,523713262 |
| F13A1 | -0,53332 | 6,2318302 | 91,1175 | 1,35E-21 | 3,12E-19 | -1,447258189 |
| SELENOP | -0,4562 | 3,1122743 | 29,00386 | 7,22E-08 | 2,84E-06 | -1,37192424 |
| HGF | -0,44038 | 1,9876509 | 13,89605 | 0,000193 | 0,00303 | -1,356964674 |
| MS4A6A | -0,42668 | 6,903627 | 131,6616 | 1,77E-30 | 9,89E-28 | -1,344141155 |
| PRR5L | -0,18533 | 2,5553107 | 3,744915 | 0,052968 | 0,214471 | -1,13707474 |
| ANGPTL4 | -0,14291 | 1,9210136 | 0,148984 | 0,699508 | 0,877811 | -1,1041264 |
| HPGD | -0,05864 | 3,2223639 | 0,455496 | 0,499737 | 0,757905 | -1,041484947 |

**D**

#### M2c Macrophages

| M1 Genes | LogFC | LogCPM | LR | P-Value | FDR | Fold change |
|---|---|---|---|---|---|---|
| CXCL11 | 3,056515 | -1,714187 | 45,36932 | 1,63E-11 | 1,69E-09 | 8,319604503 |
| CXCL10 | 2,560395 | 0,582034 | 27,54149 | 1,54E-07 | 6,83E-06 | 5,898690359 |
| ACOD1 | 2,445728 | 0,583153 | 119,1987 | 9,47E-28 | 7,46E-25 | 5,448004036 |
| IDO1 | 2,019917 | 1,462388 | 103,7551 | 2,29E-24 | 1,39E-21 | 4,05560548 |
| IFI44L | 1,839096 | 1,60344 | 164,6231 | 1,11E-37 | 1,85E-34 | 3,577857388 |
| IL1B | 1,615173 | 3,478197 | 15,30247 | 9,16E-05 | 0,001987 | 3,06348235 |
| CCL15 | 1,545454 | -1,560201 | 18,19281 | 2E-05 | 0,000533 | 2,918958104 |
| CXCL1 | 1,40164 | 1,231705 | 49,70747 | 1,78E-12 | 2,25E-10 | 2,642018353 |
| CD38 | 1,276619 | 3,115441 | 171,013 | 4,45E-39 | 8,5E-36 | 2,422705544 |
| IL12B | 1,577633 | -3,766018 | 2,903811 | 0,08837 | 0,34795 | 2,984796635 |
| **M2 Genes** | **LogFC** | **LogCPM** | **LR** | **P-Value** | **FDR** | **Fold change** |
| F13A1 | -1,30197 | 1,745999 | 68,73848 | 1,12E-16 | 2,64E-14 | -2,465646274 |
| SELENOP | -0,84189 | 3,725716 | 25,92361 | 3,55E-07 | 1,46E-05 | -1,792395723 |
| HGF | -0,74409 | 3,939544 | 15,8374 | 6,9E-05 | 0,00157 | -1,674920481 |
| MS4A6A | -0,72657 | 6,165134 | 184,6165 | 4,76E-42 | 1,27E-38 | -1,65470461 |
| TMEM37 | -0,71433 | 4,67419 | 130,9267 | 2,57E-30 | 3,12E-27 | -1,640718384 |
| PRR5L | -0,65375 | 3,796947 | 39,72088 | 2,93E-10 | 2,39E-08 | -1,573251528 |
| ANGPTL4 | -0,65299 | 0,255135 | 9,224338 | 0,002388 | 0,028861 | -1,572421318 |
| HPGD | -0,62477 | 4,922649 | 37,4014 | 9,62E-10 | 6,95E-08 | -1,541965352 |
| FAXDC2 | -0,36267 | 3,015827 | 12,19579 | 0,000479 | 0,007843 | -1,285801938 |
| HTR2B | 0,141614 | 0,523952 | 0,626865 | 0,428508 | 0,753493 | 1,103138722 |

### Supplementary Figures

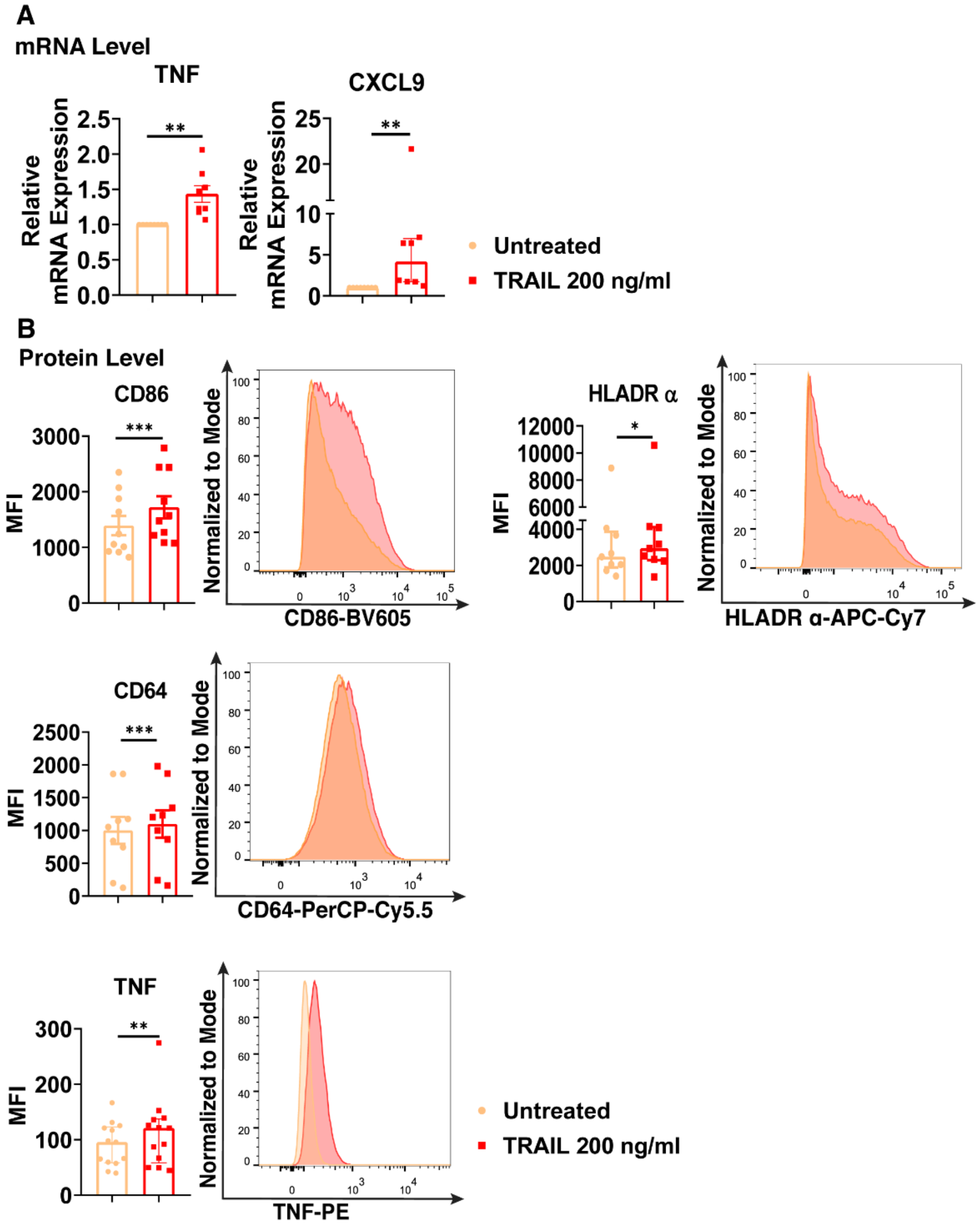


**Figure S1. TRAIL increases the expression of classical M1 markers in primary human M0 macrophages at mRNA and protein levels.** (A) M0 macrophages were stimulated with 200 ng/ml TRAIL for 8 hours. Control groups were left unstimulated. Expression of TNF and CXCL9 was analyzed by qPCR. (B) M0 macrophages were stimulated with TRAIL for 18 hours (CD86, HLA-DR alpha, CD64) or 6 hours (TNF). Control groups were left unstimulated. Expression of CD86, HLA-DR alpha, CD64 and TNF was analyzed by flow cytometry and representative plots are included. Data shown are mean ± SEM or median with interquartile range pooled from three or more independent experiments [(A) n=8, (B) n=9-13]. Statistical analyses were performed with a two-tailed paired Student's t-test or Wilcoxon matched-pairs signed-rank test between untreated and TRAIL-treated macrophages, *P<0.05, **P<0.01, ***P<0.001.

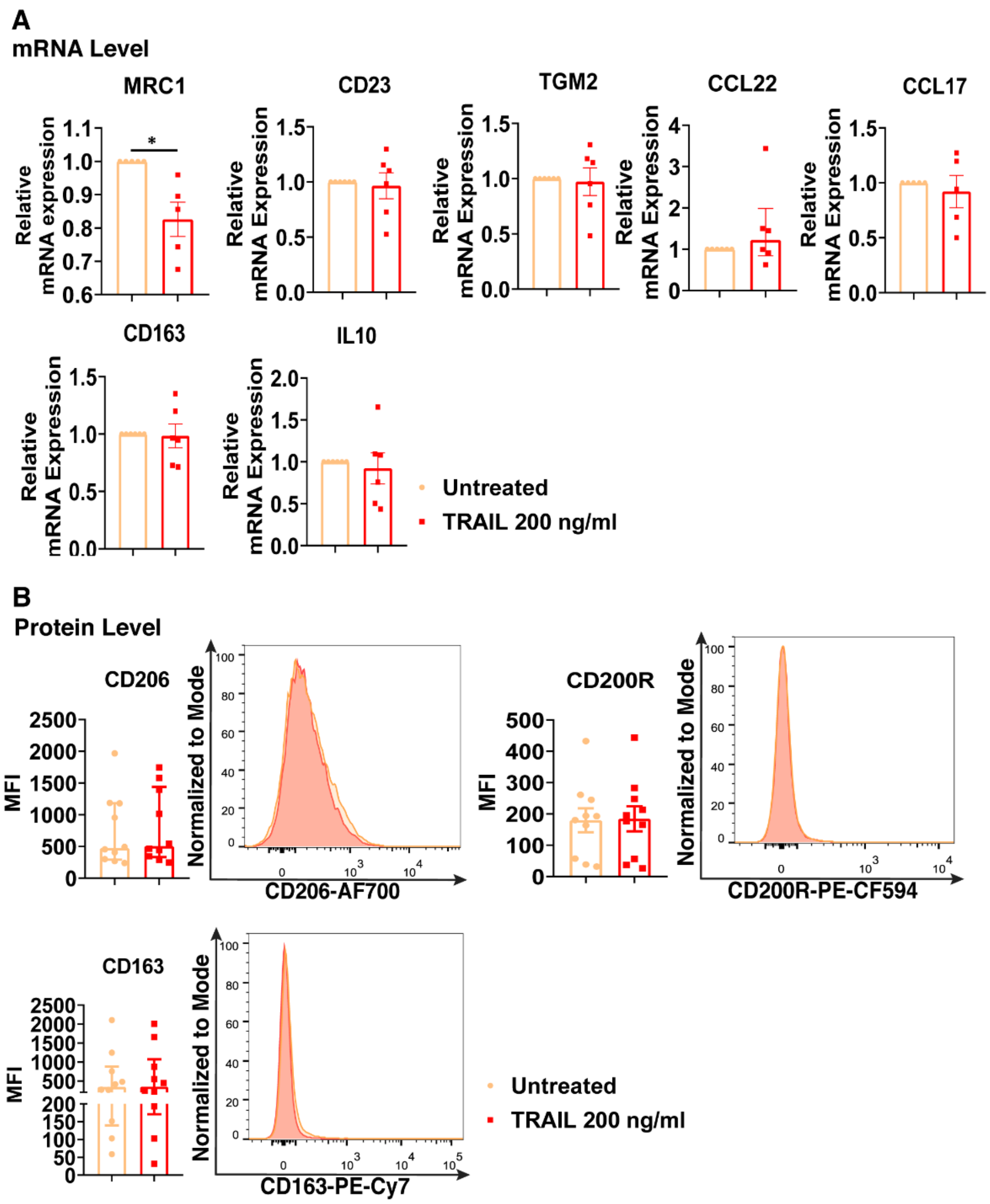


**Figure S2. TRAIL does not affect the expression classical M2 markers in primary human M0 macrophages at mRNA and protein levels.** (A) M0 macrophages were stimulated with 200 ng/ml TRAIL for 8 hours. Control groups were left unstimulated. Expression of M2 markers was analyzed by qPCR. (B) M0 macrophages were stimulated with TRAIL for 18 hours. Control groups were left unstimulated. Expression of CD206, CD200R, and CD163 was analyzed by flow cytometry, and representative plots are included. Data shown are mean ± SEM or median with interquartile range pooled from three or more independent experiments [(A) n=5-6, (B) n=10]. Statistical analyses were performed with a two-tailed paired Student's t-test or Wilcoxon matched-pairs signed-rank test between untreated and TRAIL-treated macrophages, *P<0.05.

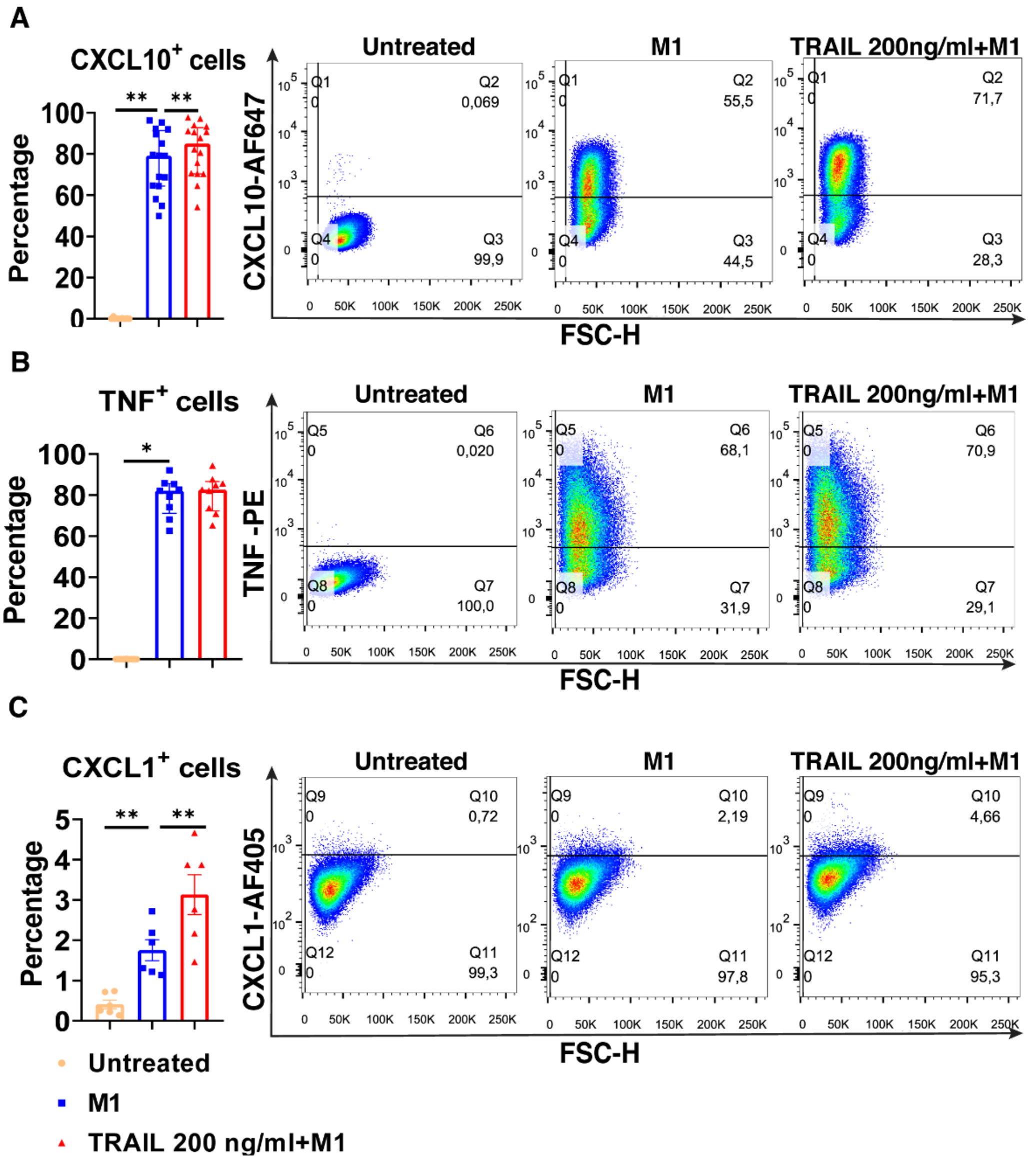


**Figure S3. TRAIL increases the percentage of M1 chemokine producing cell populations in primary human M1 macrophages at the protein level.** Macrophages were pre-stimulated with 200 ng/ml TRAIL for 2 hours and then polarized into M1 with 100 ng/ml LPS and 20 ng/ml IFNγ for 6 hours. Control groups were left unstimulated or stimulated with only M1 polarization factors for 6 hours. (A-C) The percentage of "CXCL10, TNF, and CXCL1" producing cell populations was analyzed by flow cytometry, and representative plots are shown. Data shown are mean ± SEM or median with interquartile range pooled from two or more independent experiments [(A) n=16, (B) n=9, (C) n=6]. Statistical analyses were performed with a One-way ANOVA with Sidak's multiple comparisons post-hoc test, or Friedman with Dunn's multiple comparisons post-hoc test between untreated and M1 macrophages, M1 and TRAIL-treated M1 macrophages, **P<0.01, ***P<0.001. **(Supplement of Figure 7)**

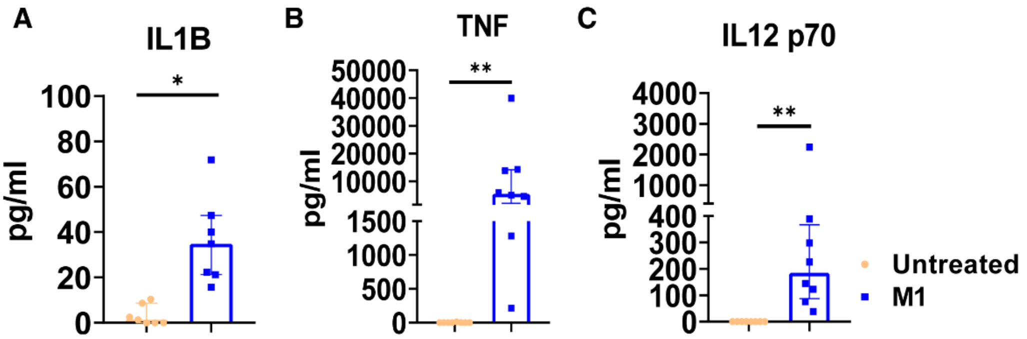


**Figure S4. M1 macrophages increases the production of IL-1β, IL-12 and TNF cytokines at the protein level.** Cytokine release of the macrophages, used in the ELISA analysis of Figure 7B, at the basal level and after M1 polarization is shown. Macrophages were polarized into M1 with 100 ng/ml LPS and 20 ng/ml IFNγ for 12 hours. Control groups were left unstimulated. Production of cytokines was analyzed by ELISA. Data shown are median with interquartile range pooled from three or more independent experiments [(A) n=7, (B) n=8, (C) n=8]. Statistical analyses were performed with a Wilcoxon matched-pairs signed-rank test between untreated and M1 macrophages, $*P<0.05$, $**P<0.01$ **(Supplement of Figure 7).**

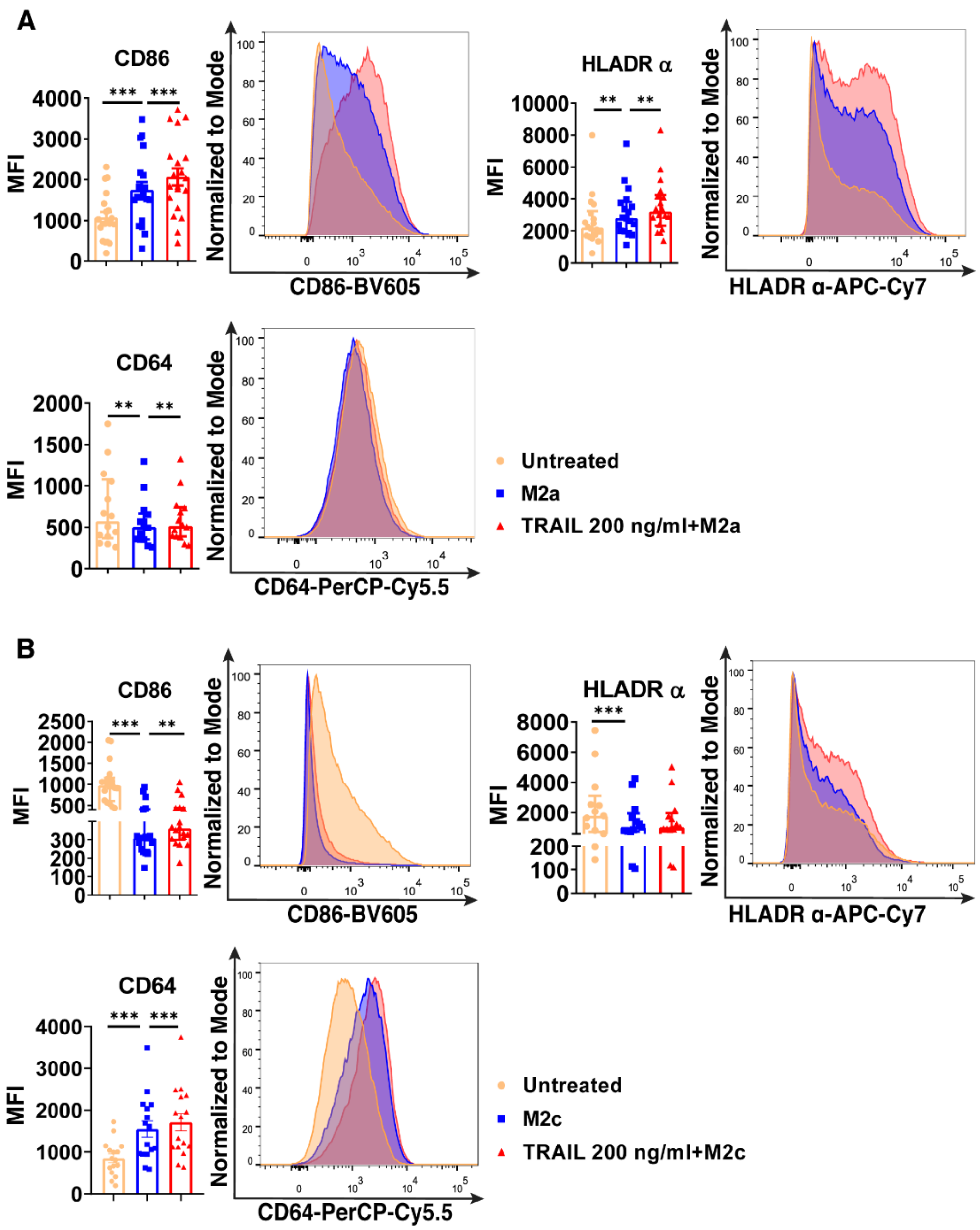


**Figure S5. TRAIL increases the expression of classical M1 markers in primary human M2a and M2c macrophage subtypes at the protein level.** Macrophages were pre-stimulated with 200 ng/ml TRAIL for 6 hours and then polarized into M2a with 20 ng/ml IL-4 or M2c with 10 ng/ml IL-10 for 12 hours. Control groups were left unstimulated or stimulated with only M2a or M2c polarization factors for 12 hours. Expression of cell surface M1 markers was analyzed by flow cytometry in (A) M2a and (B) M2c macrophages and representative plots are included. Data shown are mean ± SEM or median with interquartile range pooled from four or more independent experiments [(A) n=14-20, (B) n=13-18]. Statistical analyses were performed with a One-way ANOVA with Sidak's multiple comparisons post-hoc test, or Friedman with Dunn's multiple comparisons post-hoc test between untreated and M2 macrophages, M2 and TRAIL-treated M2 macrophages, *P<0.05, **P<0.01, ***P<0.001. **(Supplement of Figure 8).**

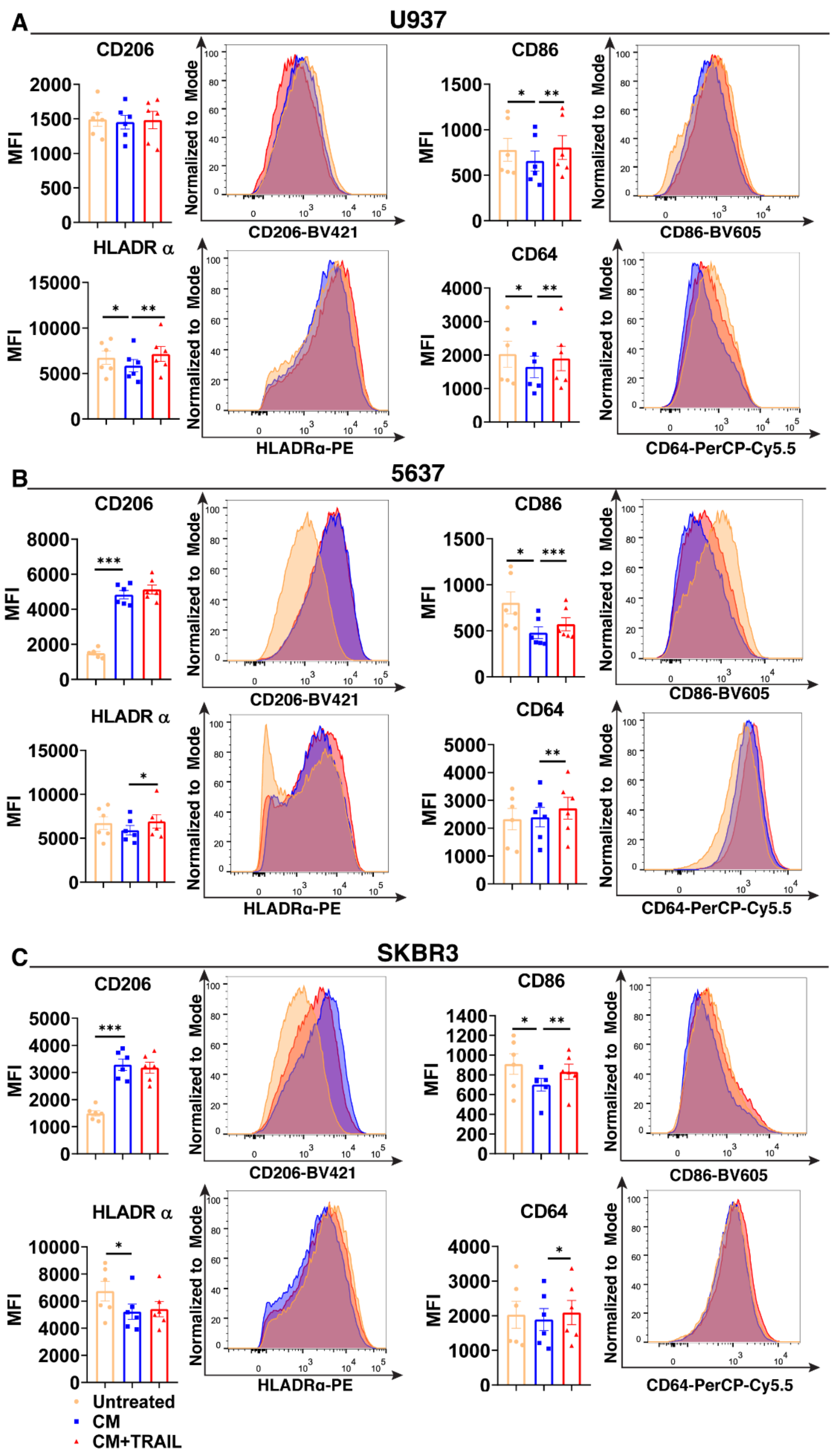


**Figure S6. TRAIL increases the expression of M1 markers in tumor-associated macrophages (TAMs).** Macrophages were polarized into TAMs with conditioned media (CM) of (A) U937, (B) 5637, and (C) SKBR3 tumor cell lines for 72 hours. TAMs were stimulated with 200 ng/ml TRAIL in the last 18 hours of incubation without removing the tumor CM. Control group macrophages were left unstimulated or only treated with tumor CM for 72 hours. Expression of CD206 M2 marker and CD86, HLADRα and CD64 M1 markers was analyzed by flow cytometry and representative plots are included. Data shown are mean ± SEM pooled from two independent experiments (n=6). Statistical analyses were performed with a One-way ANOVA with Sidak's multiple comparisons post-hoc test between untreated macrophages and tumor CM treated macrophages (TAMs), TAMs and TRAIL treated-TAMs, $^{*}P<0.05$, $^{**}P<0.01$, $^{***}P<0.001$.

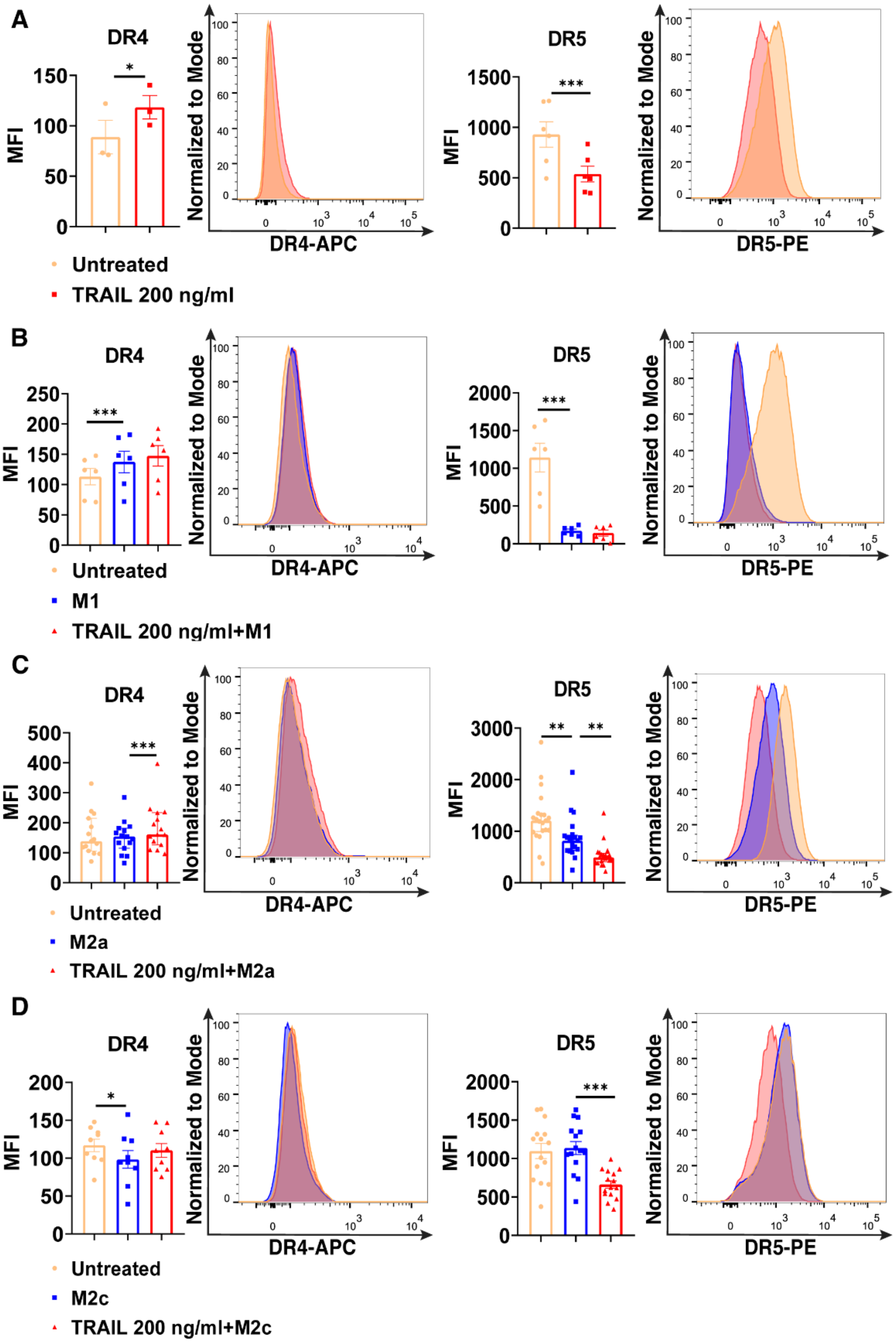


**Figure S7. TRAIL increases DR4 and decreases DR5 expression at the cell surface in primary human macrophage subtypes.** Macrophages were stimulated with 200 ng/ml TRAIL for 18 hours or pre-stimulated with TRAIL for 6 hours and then polarized into M1 with 100 ng/ml LPS and 20 ng/ml IFNγ, M2a with 20 ng/ml IL-4, or M2c with 10 ng/ml IL-10 for 12 hours. Control groups were left unstimulated or stimulated with only M1, M2a, or M2c polarization factors for 12 hours. Expression of DR4 and DR5 in (A) M0, (B) M1, (C) M2a, (D), and M2c macrophages was analyzed by flow cytometry, and representative plots are included. Data shown are mean ± SEM or median with interquartile range pooled from three or more experiments [(A) n=3-6, (B) n=6, (C) n=15-21, (D) n=9-15]. Statistical analyses were performed with a two-tailed paired Student's t-test, One-way ANOVA with Sidak's multiple comparisons post-hoc test, or Friedman with Dunn's multiple comparisons post-hoc test between untreated and polarized macrophages, untreated/polarized and TRAIL-treated macrophages, *P<0.05, ***P<0.001.

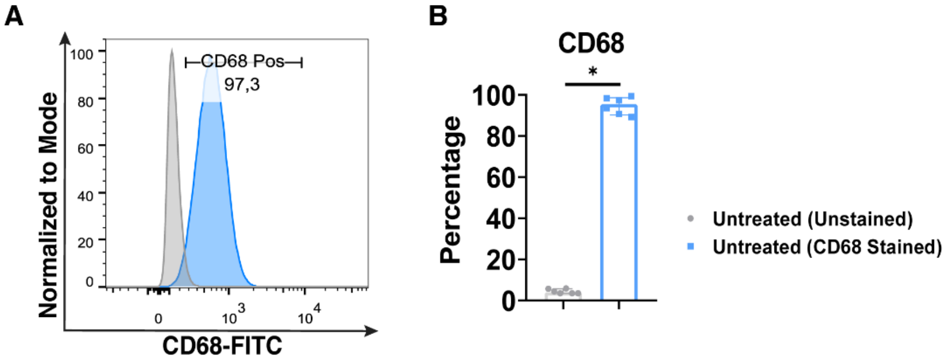


**Figure S8. Primary human monocytes are successfully differentiated into macrophages *in vitro*.** Monocytes isolated from PBMCs were differentiated into macrophages by incubating the cells in 10 ng/ml M-CSF containing R5 media for 7 days. Expression of human macrophage maturation marker CD68 in M0 macrophages was analyzed by flow cytometry. (A) Representative plot and (B) Bar graph show the percentage of $CD68^+$ cells. Data shown are median with interquartile range pooled from three independent experiments (n=6). Statistical analyses were performed with a Wilcoxon matched-pairs signed-rank test, between unstained and CD68 stained groups, *P<0.05.
